# Behavioral origin of sound-evoked activity in mouse visual cortex

**DOI:** 10.1101/2021.07.01.450721

**Authors:** Célian Bimbard, Timothy PH Sit, Anna Lebedeva, Charu B Reddy, Kenneth D Harris, Matteo Carandini

## Abstract

Sensory cortices can be affected by stimuli of multiple modalities and are thus increasingly thought to be multisensory. For instance, primary visual cortex (V1) is influenced not only by images but also by sounds. Here we show that the activity evoked by sounds in V1 is highly stereotyped across neurons and even across mice. It resembles activity measured elsewhere in the brain and is independent of projections from auditory cortex. Its low-dimensional nature starkly contrasts the high dimensional code that V1 uses to represent images. Furthermore, this sound-evoked activity can be precisely predicted by small body movements that are elicited by each sound and are highly stereotyped across trials and across mice. Thus, neural activity that is apparently multisensory may simply arise from low-dimensional signals associated with changes in internal state and behavior.

## Introduction

Many studies suggest that all cortical sensory areas, including primary ones, are multisensory^1^. For instance, mouse primary visual cortex (V1) is influenced by sounds. Sounds may provide V1 with global inhibition^2^, modify the neurons’ orientation tuning^3,4^, boost detection of visual events^5^, or even provide tone-specific information, reinforced by prolonged exposure^6^ or training^7^. This sound-evoked activity is thought to originate from direct projections from the auditory cortex^2,3,5,7^: it can be suppressed by inhibition of the auditory cortex^2,5^, and it may be mimicked by stimulation of auditory fibers^2,3^.

Here, we consider a possible alternative explanation for these multisensory signals, based on low-dimensional changes in internal state and behavior^8^. Behavioral and state signals have profound effects on sensory areas. For instance, the activity of V1 neurons is powerfully affected by signals related to running^9,10^, pupil dilation^10,11^, whisking^12^, and other movements^13^. These behavioral and state signals are low-dimensional and largely orthogonal^12^ to the high-dimensional code that V1 uses to represent images^14^.

Specifically, we conjecture that the activity evoked by sounds in V1 may reflect sound-elicited changes in internal state and behavior. This seems possible, because sounds can change internal state and evoke uninstructed body movements^15-19^. We thus asked whether sound-evoked activity in V1 may have the typical attributes of behavioral signals: low dimension^12^ and a broad footprint^13,20,21^ that extends to brain regions beyond the cortex^12^. Moreover, we reasoned that if sound-evoked activity is related to behavioral and state signals, it should be independent of direct inputs from auditory cortex and it should be predictable from the behavioral effects of sounds.

To test these predictions, we recorded the responses of hundreds of neurons in mouse V1 to audiovisual stimuli, while filming the mouse to assess the movements elicited by the sounds. As predicted by our hypothesis, the activity evoked by sounds in V1 had a low dimension: it was largely one-dimensional. Moreover, it was essentially identical to activity evoked in another brain region: the hippocampal formation. Furthermore, it was independent of direct projections from auditory cortex, and it tightly correlated with the uninstructed movements evoked by the sounds. These movements were small but specific to each of the sounds and highly stereotyped across trials and across mice. Thus, a large fraction of the multisensory activity that has been observed in visual cortex may have a simpler, behavioral origin.

## Results

To explore the influence of sounds on V1 activity, we implanted Neuropixels 1.0 and 2.0 probes^22,23^ in 8 mice, and recorded during head fixation while playing naturalistic audiovisual stimuli. We selected 11 naturalistic movie clips^24^, each made of a video (gray-scaled) and a sound (loudness 50-80 dB SPL, Suppl. Figure 1), together with a blank movie (gray screen, no sound). On each trial, we presented a combination of the sound from one clip and the video from another (144 combinations repeated 4 times, in random order).

### Sounds evoke stereotyped responses in visual cortex

We then identified the visual and auditory components of each neuron’s sensory response. A typical V1 neuron responded differently to different combinations of videos and sounds (Figure 1a). To characterize these responses, we used a marginalization procedure similar to factorial ANOVA. To measure the neuron’s video-related responses (Figure 1b) we computed its mean response to each video, averaged across all concurrent sounds. Similarly, to characterize the neuron’s sound-related responses (Figure 1c) we computed the mean response to each sound, averaged across all concurrent videos. These measures were ‘marginalized’ by subtracting the grand average over all videos and sounds (Figure 1d).

**Figure 1.**
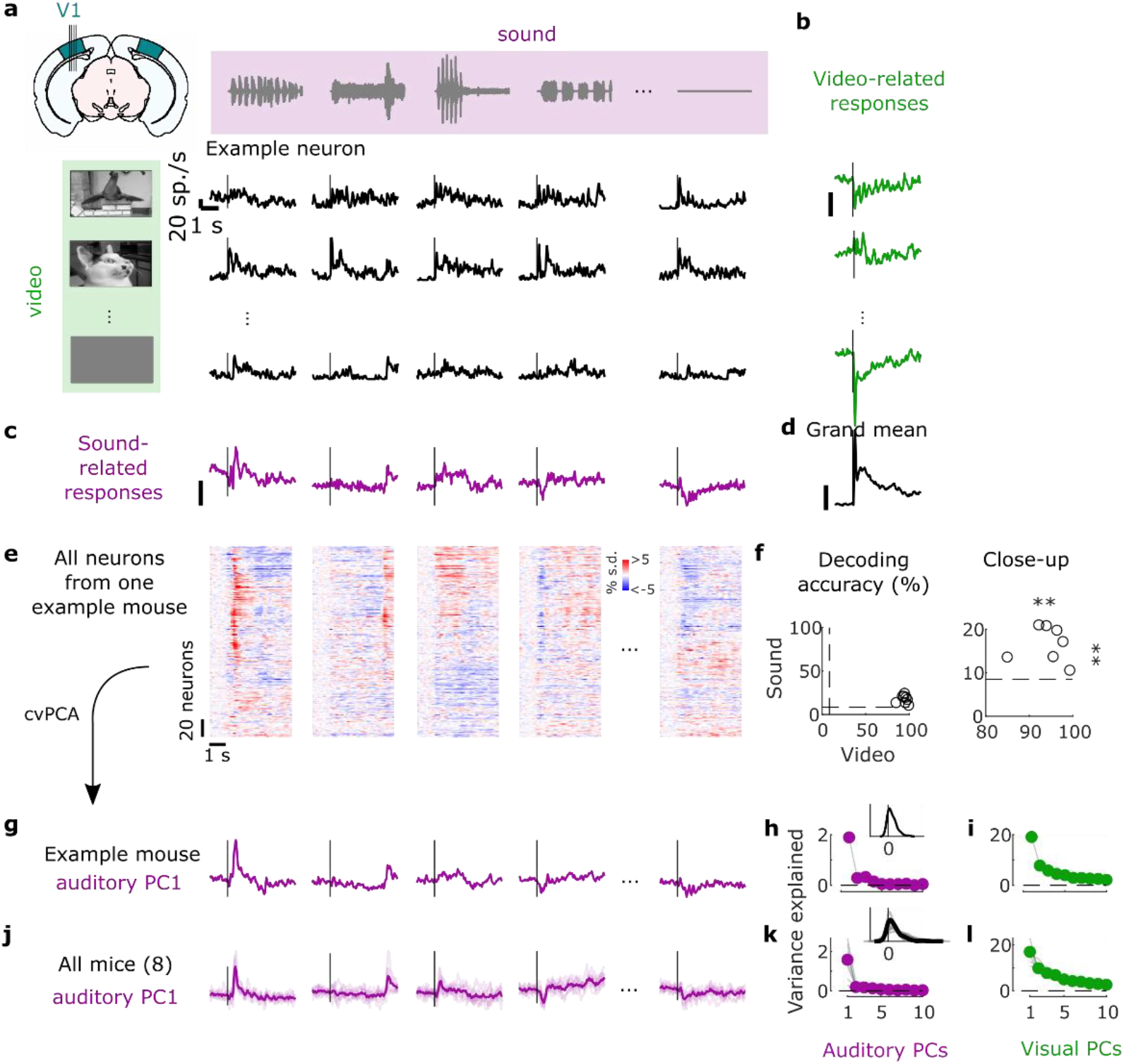
Sounds evoke stereotyped responses in visual cortex. **a.** Responses of an example neuron to combinations of sounds (columns) and videos (rows). Curves show the average response over 4 repeats. **b.** Video-related time courses (averaged over all repeats and sound conditions, relative to grand average) for the example neuron in **a. c.** Same, for the sound-related time courses. **d.** Grand average over all conditions for the neuron. Scale bars as in **b-d**: 20 spikes/s). **e.** Sound-related time courses for all 212 neurons in one experiment, sorted using *rastermap*^12^ **f.** Decoding accuracy for video vs. sound for all 8 mice (**: p-value<0.01). Dashed lines show chance level (1/12). **g.** Time courses of the first principal component of the sound-related responses in **e** (‘auditory PC1’, arbitrary units), obtained through cross-validated principal components analysis (cvPCA). **h.** Fraction of total variance explained by successive auditory PCs, for this example mouse; inset: distribution of the weights of auditory PC1 (arbitrary units), showing an overall positive distribution. **i**. Same, for visual PCs. **j-,l** Same as **g-I**, averaged across mice (thin lines denote individual mice).

Sounds evoked activity in a large fraction of V1 neurons, and this activity was reliably different across sounds. Some sounds barely evoked any activity, while others evoked stereotyped responses, at different points in time (Figure 1e). From the marginalized single-trial population responses, we could significantly decode not only the identity of each video (with 94 ± 2% accuracy, s.e., p = 0.0039, right-tailed Wilcoxon sign rank test, n = 8 mice) but also the identity of each sound (with 18 ± 2% accuracy, p = 0.0039, right-tailed Wilcoxon sign rank test, n = 8 mice, Figure 1f).

The activity evoked by sounds was highly stereotyped across responsive neurons, to the point that it was close to one-dimensional. We applied cross-validated Principal Component Analysis^14^ (cvPCA) to the marginalized population responses, and found that the time course of the first dimension (“auditory PC1”) for each sound was similar to the responses evoked in individual neurons and different across sounds (Figure 1g). This first dimension explained most (55%) of the cross-validated sound-related variance (1.9% of the total variance) with subsequent dimensions explaining much smaller fractions (Figure 1h). Furthermore, neurons showed distributed yet overall positive weights on this first PC, indicating a largely excitatory effect of sound (Figure 1h, inset). Thus, in the rest of the paper we will illustrate sound-evoked activity by using the time course of this single “auditory PC1”.

Similar results held across mice: the activity evoked by sounds in V1 was largely one-dimensional (auditory PC1 explained 54 ± 3% of the sound-related variance, s.e., n = 8 mice), and the first principal component across mice had similar time courses and similar dependence on sound identity (Figure 1j,k). Indeed, the correlation of auditory PC1 timecourses evoked in different mice was 0.35, close to the test-retest correlation of 0.43 measured within individual mice. Again, in all mice, the neuron’s weights for the auditory PC1 were widely distributed, with a positive bias (p = 0.0078, two-tailed Wilcoxon sign rank test on the mean, n = 8 mice, Figure 1k). Thus, sounds evoke essentially one-dimensional population activity, which follows a similar time course even across brains.

In contrast, the activity evoked by videos in V1 neurons was markedly larger and higher-dimensional. The first visual PC explained a much higher fraction of total variance than the first auditory PC (17 ± 1% vs 1.6% ± 0.3, s.e., n = 8 mice; Figure 1i,l). Furthermore, higher visual PCs explained substantial amounts of variance, as previously reported^14^, unlike higher auditory PCs.

### Sounds evoke stereotyped responses in hippocampal formation

We next investigated whether these auditory-evoked signals were specific to visual cortex. Thanks to the length of Neuropixels probes, while recording from V1 we simultaneously recorded from the hippocampal formation (dorsal and pro-subiculum, dentate gyrus, and CA3, Figure 2). Inspection of the Allen Mouse Brain Connectivity Atlas^25^ indicates that these regions receive little input from auditory cortex and auditory thalamus.

**Figure 2.**
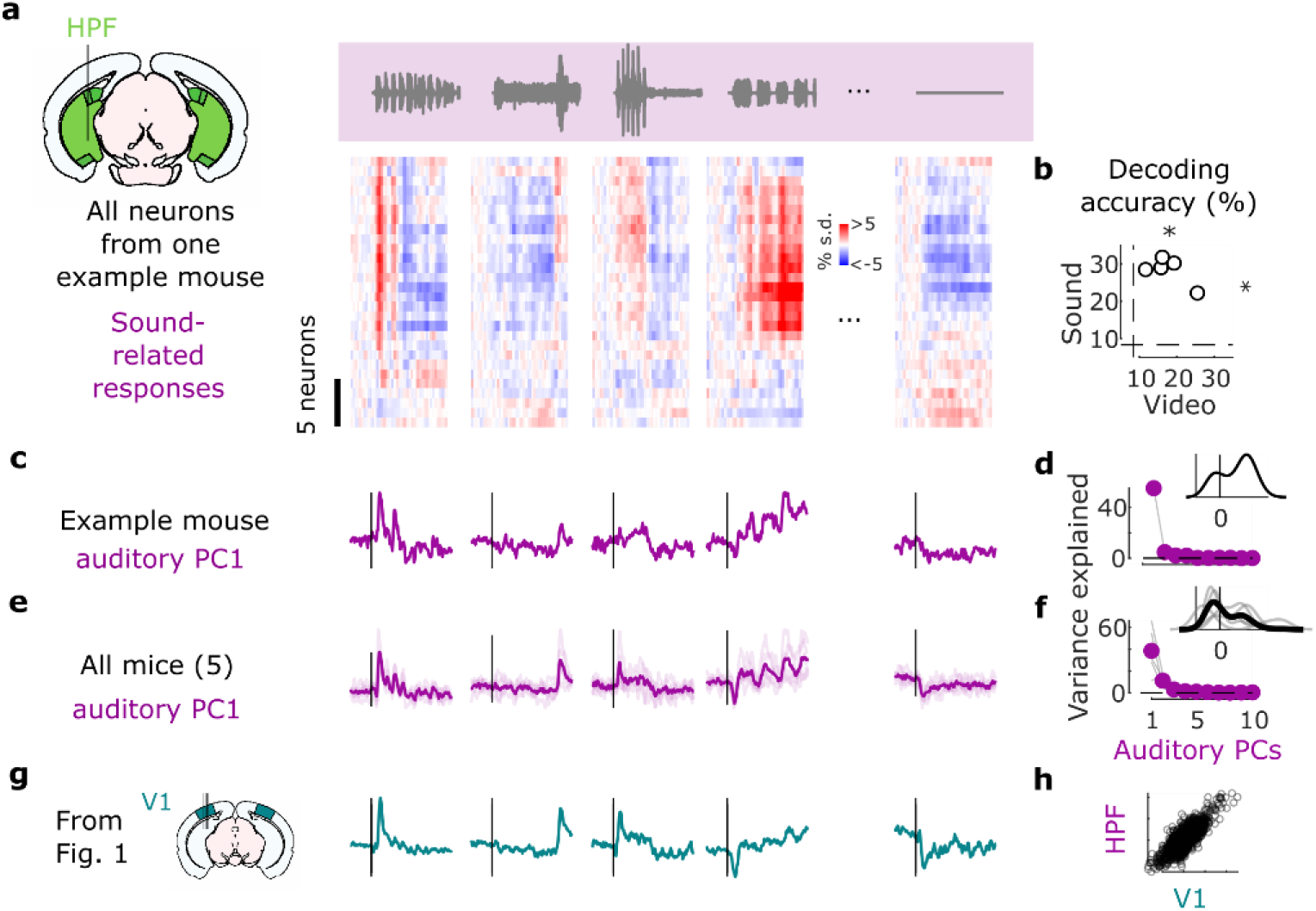
Sounds evoke stereotyped activity in hippocampal formation. **a.** Sound-related time courses for all 28 neurons in in hippocampal formation (HPF) in one experiment, sorted using *rastermap*^12^ **b.** Decoding accuracy for video vs. sound for all 8 mice (*: p-value<0.05). Dashed lines show chance level (1/12). **c.** Time courses of the first principal component of the sound-related responses in **a** (‘auditory PC1’, arbitrary units), obtained through cross-validated principal components analysis (cvPCA). **d.** Fraction of total variance explained by successive auditory PCs, for this example mouse; inset: distribution of the weights of auditory PC1 (arbitrary units). **e,f.** Same as **c,d** for all mice (individual mice as thin lines). **g.** Time courses of the auditory PC1 in visual cortex (taken from Figure 1) for comparison. **h.** Comparison of the auditory PC1 from HPF (taken from **e**) and from V1 (taken from Figure 1); arbitrary units.

Neurons in the hippocampal formation exhibited strong sound-evoked activity. Sounds evoked activity that was largely similar across cells and different across sounds (Figure 2a). As in visual cortex, this activity in single trials could be used to decode sound identity (28 ± 2% and to a lesser extent video identity 18 ± 2%, p = 0.031 for both, two-tailed Wilcoxon sign rank test, n = 5 mice; Figure 2b). Projection of the sound-related activity along the auditory PC1 showed different time courses across sounds (Figure 2c,e), and this first PC explained most of the sound-related variance (64 ± 12% Figure 2d,f). Similarly, the representation of videos was also low-dimensional (Suppl. Figure 2b). Overall, the time courses of auditory PC1 were very similar to those observed in visual cortex: their correlation was r = 0.82 (Figure 2g,h).

The activity evoked by sounds in the hippocampal formation was remarkably similar to the activity evoked in visual cortex. Indeed, the time courses of the auditory PC1 in the two regions, averaged over mice, were hardly distinguishable (compare Figure 2e to Figure 2g), with a correlation of r = 0.82 (Figure 2h).

### Sound-evoked activity in visual cortex is independent of inputs from auditory cortex

We next returned to the activity evoked by sounds in visual cortex and asked if this activity is due to projections from auditory cortex, as has been proposed^2,3,5,7^. We performed transectomies^2^ to cut the fibers between auditory and visual areas in one hemisphere and recorded bilaterally while repeating our audiovisual protocol (Figure 3a). The cut ran along the whole boundary between auditory and visual areas and was deep enough to reach into the white matter (Suppl. Figure 3a-c). We used the software packages *brainreg*^26-28^ to quantify the cuts and *brainrender*^29^ to visualize their extent in 3D (Figure 3b, Suppl. Figure 3d). To estimate the fraction of fibers from auditory to visual areas that were cut, we extracted the relevant trajectories of fibers from the Allen Mouse Brain Connectivity Atlas^25^, and intersected it with the location of our cut. We thus estimated that the auditory input to the visual areas ipsilateral to the cut would be decreased by a factor >3.5 on average compared to the contralateral side (4.7, 2.4 and 3.6 for each mouse, Figure 3c, Suppl. Figure 3e-g). Thus, if auditory evoked activity in visual cortex originates from auditory cortex, it should be drastically reduced on the cut side.

**Figure 3.**
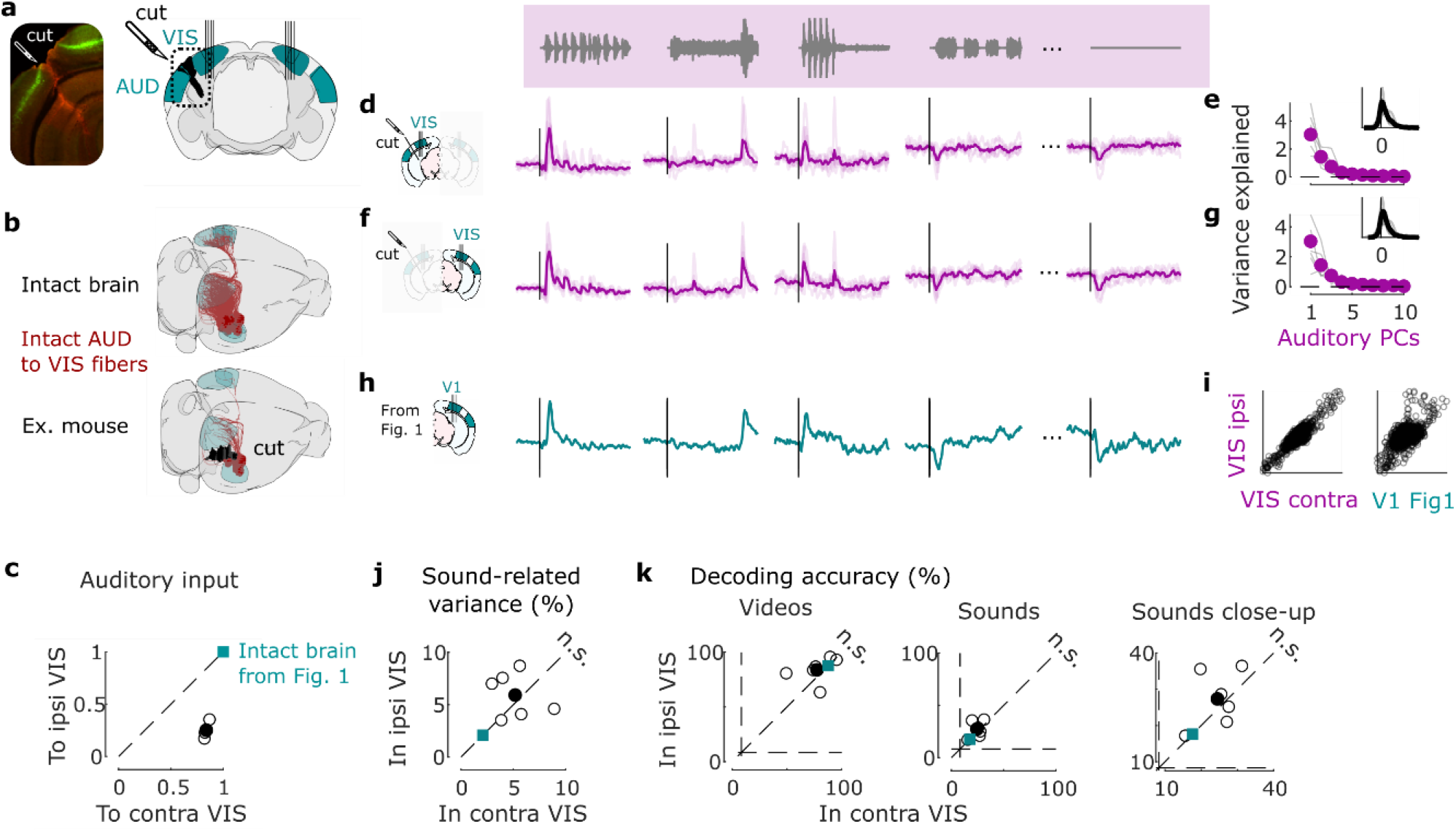
Sound-evoked activity in visual cortex is independent of projections from auditory cortex. **a.** Coronal views of a transectomy cutting the connections between auditory (AUD) and visual (VIS) cortex in one hemisphere, showing histology *left*) and reconstruction of the cut using brainrender (*right*). After the cut, bilateral recordings are performed in visual cortex. **b**. 3D visualizations showing AUD to VIS fibers (*red*) in an intact brain (*top*) vs. after the cut in one example mouse (*bottom*). **c**. Auditory input to the sides contralateral vs. ipsilateral to the cut for all 3 mice (open dots) and their average (filled dot), normalized by the input expected in intact brains (turquoise dot). **d**. Time courses of the first principal component of the sound-related responses (‘auditory PC1’) on the side ipsilateral to the cut (average over all mice). Thin curves lines show individual mice. **e.** Fraction of total variance explained by successive auditory PCs on the side ipsilateral to the cut; inset: distribution of the weights of auditory PC1 for all mice. **f,g.** Same as **d,e** for the side contralateral to the cut. **h.** Time courses of the auditory PC1 in visual cortex of intact, control mice (taken from Figure 1) for comparison. **i.** Comparison of the auditory PC1 from the sides contralateral and ipsilateral to the cut (*left*, taken from **d** vs. **f**) and from V1 (right, taken from **b** vs. Figure 1); all arbitrary units. **j**. Sound-related variance explained by the first 4 auditory PCs on the ipsi- vs. contra-lateral side, showing individual sessions (open dots), their average (black dot), and the average across control mice (turquoise dot). There is no significant deviation from the diagonal. **k**. Decoding accuracy for videos (*left*) and sounds (*middle* and *right*, showing close-up). Symbols as in **j**. There is no significant deviation from the diagonal, i.e., no significant difference between the cut and uncut sides.

The activity evoked by sounds in visual cortex was similar on the cut and the uncut side. Indeed, the time course of the activity projected along auditory PC1 on the side of the cut (Figure 3d,e) was essentially identical to the time course of auditory PC1 in the opposite hemisphere (r = 0.9, Figure 3f,g) and barely distinguishable from the one measured in the control mice (cut: r = 0.62 / uncut: r = 0.56, Figure 3h,i). The distribution of the variance explained by the first auditory PCs and the distribution of neuronal weights on the auditory PC1 were similar in the two sides (Figure 3e vs. g). The total variance of the activity related to sounds on the cut side was on average equal to the sound-related variance on the uncut side (Figure 3j, see Suppl. Figure 2c for all eigenspectra) and was significantly larger than expected from the few auditory fibers that were spared by the transectomies (p = 0.031, two-tailed paired sign rank test, n = 6 experiments across 3 mice). Furthermore, decoding accuracy was similar across sides for both sounds (cut: 27 ± 3% / uncut: 25 ± 2%, p = 0.016 for both, right-tailed Wilcoxon sign rank test; comparison: p = 0.56, two-sided paired Wilcoxon sign rank test) and videos (cut: 87 ± 6%/ uncut: 81 ± 6%, p = 0.016 for both, right-tailed Wilcoxon sign rank test; comparison: p = 0.43, two-sided paired Wilcoxon sign rank test; Figure 3k).

These results indicate that the activity evoked by sounds in visual cortex cannot be explained by inputs from auditory cortex.

### Sounds evoke stereotyped, uninstructed behaviors

Sounds evoked uninstructed body movements that were small but highly stereotyped across trials and across mice, and different across sounds. To measure body movements, we used a wide-angle camera that imaged the head, front paws, and back of the mice (Figure 4a). Sounds evoked a variety of uninstructed movements, ranging from rapid startle-like responses <50 ms after sound onset to more complex, gradual movements (Figure 4b, see Suppl. Figure 4 for all sounds). These movements were remarkably similar across trials and mice. The main and most common type of sound-evoked movements were subtle whisker twitches (Suppl. Video 1), which we quantified by plotting the first principal component of facial motion energy^12^ (Figure 4b). These movements were influenced by sound loudness, and to some extent by frequency, but not by spatial location (Suppl. Figure 5). Moreover, sounds evoked stereotyped changes in arousal, as observed by the time courses of pupil size, which were highly consistent across trials and mice (Suppl. Figure 6).

**Figure 4.**
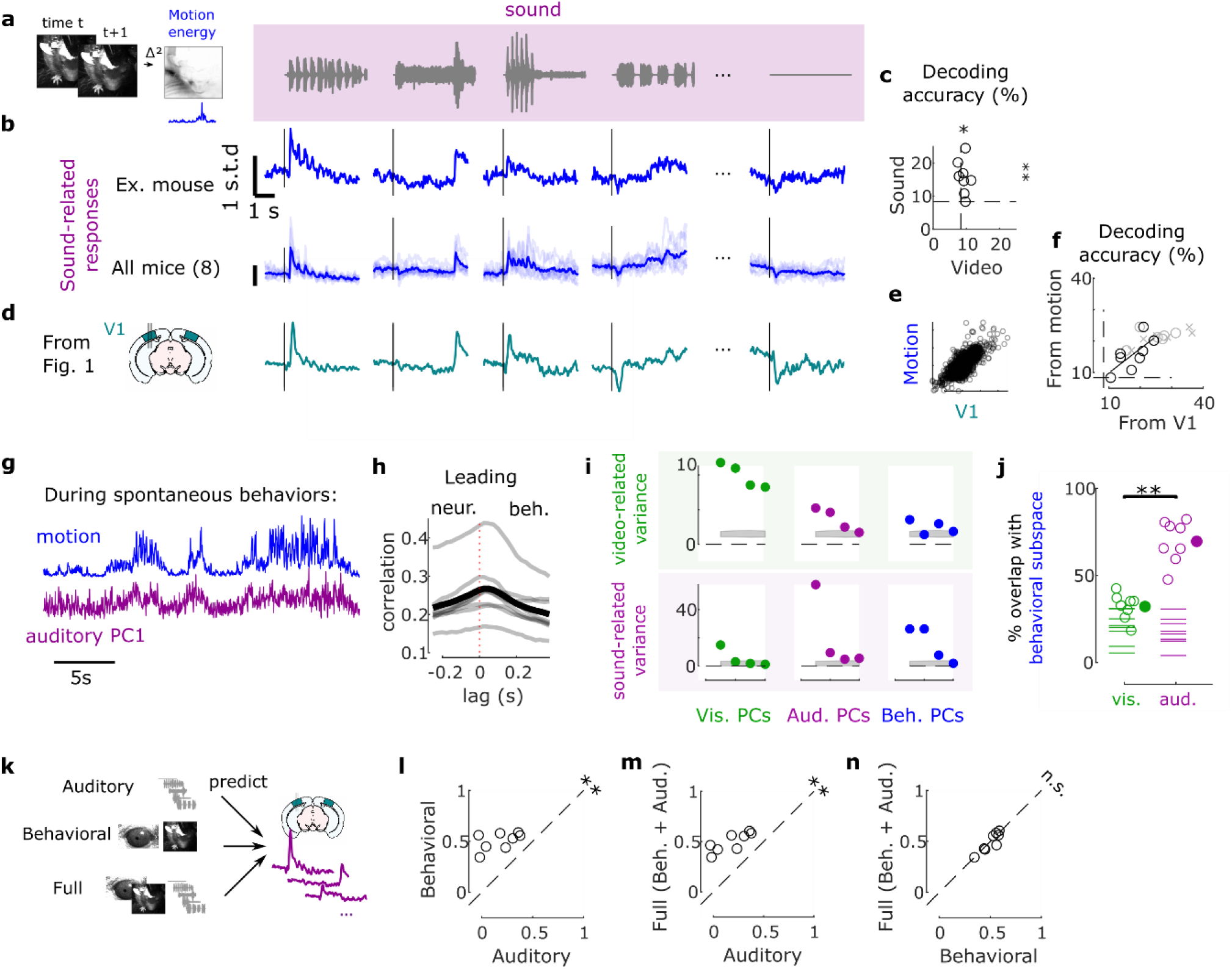
Sounds evoke stereotyped, uninstructed behavioral responses that explain neural activity. **a.** Extraction of motion PCs from videos of the mouse face. **b.** Sounds evoked changes in the first motion PC, both in an example mouse (*top*) and all mice (*bottom*). Scale bar: 1 s.d. **c.** Decoding of video and sound identity from the first 128 motion PCs was significantly above chance level (dashed lines) (*: p-value<0.05, **: p-value<0.01). **d.** Time courses of the auditory PC1 in visual cortex (from Figure 1). **e.** Comparison of the time courses of motion (taken from **b**) and of the auditory PC1 from V1 (taken from Figure 1); all arbitrary units. **f.** Across mice, there was a strong correlation between the accuracy of sound decoding from facial motion and from V1 activity. The linear regression is performed on the control mice from Figure 1 (black dots). Data from transectomy mice (gray dots) confirm the trend, both in the cut side (crosses) and on the uncut side (circles). **g.** Time course of facial motion (*top*) and of V1 activity along auditory PC1 (*bottom*) in the absence of any stimulus, for an example mouse. **h**. Cross-correlogram of these time courses, for individual mice (*gray*) and their average (*black*). The positive lag indicates that movement precedes neural activity. **i.** Video- and sound-related variance explained by neural activity along the visual (*left*), auditory (*middle*), or behavioral (*right*) subspaces (first 4 PCs of each subspace), for one example mouse. The gray regions show 90% confidence intervals expected by chance (random components). **j.** Overlap between the auditory or the visual subspace and the behavioral subspace for each mouse (open dots) and all mice (filled dot). Dashed lines show the significance threshold (95^th^ percentile of the overlap with random dimensions) for each mouse. **k.** Schematics of the 3 encoding models trained to predict the average sound-related activity in the auditory subspace. **l-n.** Cross-validated correlation of the actual sound responses and their predictions for all mice, comparing different models (Auditory: sounds only, Behavioral: eye and body movements only, Full: all predictors; **: p-value<0.01).

Because sound-evoked movements were different across sounds and similar across trials, we could use them to decode sound identity with 16 ± 2% accuracy (s.e., p = 0.0039, right-tailed Wilcoxon sign rank test, n = 8 mice, Figure 4c). This accuracy was not statistically different from the 18 ± 2% accuracy of sound decoding from neural activity in visual cortex (p = 0.20, two-sided paired Wilcoxon sign rank test), suggesting a similar level of single-trial reliability in behavior and in neural activity.

### Sound-evoked behaviors predict sound-evoked responses in visual cortex

The body movements evoked by sounds had a remarkably similar time course to the activity evoked by sounds in area V1 (Figure 4b,d). The two were highly correlated across time and sounds (r = 0.75, Figure 4e, see Suppl. Figure 4 for all sound-related timecourses). Furthermore, the accuracy of decoding sound identity from V1 activity and from behavior was highly correlated across mice (r = 0.71, p = 0.048, F-statistic vs. constant model, n = 8 mice, Figure 4f), suggesting that sound-specific neural activity was higher in mice that moved more consistently in response to sounds. As it happens, the cohort of transectomy mice showed higher sound decoding accuracy from their behavior compared to the main cohort. Consistent with our hypothesis, these same mice showed higher sound decoding accuracy from their V1 activity, regardless of hemisphere. Finally, the neural activity along auditory PC1 correlated with movements even during spontaneous behavior, when no stimulus was presented (Pearson correlation 0.27 ± 0.03, s.e., Figure 4g). Movement preceded neural activity by a few tens of milliseconds (28 ± 7ms, s.e., p = 0.031, two-sided Wilcoxon sign rank test, n = 8 mice, Figure 4h, see Suppl. Figure 7 for the hippocampal formation and for both sides of the visual cortex in transectomy experiments).

Another similarity between the neural activity evoked by sounds and by movement could be seen in their subspaces^12^, which substantially overlapped with each other. To define the behavioral subspace, we applied reduced-rank regression to predict neural activity from movements during the spontaneous period (in the absence of stimuli). This behavioral subspace largely overlapped with the auditory subspace: the first 4 components of the movement-related subspace explained 69 ± 4% (s.e., p < 0.05 for all mice separately, randomization test) of the sound-related variance, much more than the video-related variance^12^ (32 ± 3%, comparison: p = 0.0078, two-sided paired Wilcoxon sign rank test, Figure 4I,j). We observed a similar overlap in the hippocampal formation, and on both sides of visual cortex in the transectomy experiments (Suppl. Figure 7).

We then asked to what extent body movements could predict sound-evoked neural activity in V1, and to this end we compared three models (Figure 4k, Suppl. Figure 8): (1) a purely *auditory* model where the time course of neural activity depends only on sound identity (equivalent to a test-retest); (2) a purely *behavioral* model where neural activity is predicted by pupil area, eye position/motion, and facial movements; (3) a *full* model where activity is due to the sum of both factors.

We fitted the models on the marginalized (i.e., sound-related), single-trial data and used them to predict trial-averages of sound-evoked activity (rather than trial to trial fluctuations). The auditory model is thus expected to perform well regardless of the origin of sound-evoked activity, since it is equivalent to a test-retest prediction, which would fit perfectly with an infinite number of trials. By contrast, the behavioral model will perform well only if behavioral variables do predict the trial-averaged sound-evoked responses.

This analysis revealed that the sounds themselves were unnecessary to predict sound-evoked activity in visual cortex: the body movements elicited by sounds were sufficient. As expected, the auditory model was able to capture much of this activity. However, it performed worse than the full model and the behavioral models (p = 0.0078, two-sided paired Wilcoxon sign rank test, Figure 4l,m). These models captured not only the average responses to the sounds, but also the fine differences in neural activity between the train and test set, which the auditory model cannot predict (because the two sets share the same sounds). Remarkably, the behavioral model performed just as well as the full model (p = 0.74, two-sided paired Wilcoxon sign rank test, Figure 4n), indicating that the extra predictors - the sounds themselves - were unnecessary to predict sound-evoked activity. Further analysis indicated that the main behavioral correlates of sound-evoked activity in V1 were movements of the body and especially of the whiskers, rather than the eyes (Suppl. Figure 9).

By contrast, and indeed as expected for a brain region that encodes images, a purely visual model explained a high fraction of the neural activity evoked by videos (Suppl. Figure 10a-c, j-o). In the hippocampal formation, finally, the behavioral model explained both the sound- and video-evoked activity, suggesting that any visual or auditory activity observed there is largely related to movements (Suppl. Figure 10d-i).

Further confirming the role of body movements, we found that trial-by-trial variations in sound-evoked V1 activity were well-predicted by trial-by-trial variations in body movement (Suppl. Figure 11). The movements elicited by each sound were stereotyped but not identical across trials. The behavioral model and the full model captured these trial-by-trial variations, which could not be captured by the auditory model because (by definition) the sounds did not vary across trials. The trial-by-trial variations of the visual cortex’s auditory PC1 showed a correlation of 0.39 with its cross-validated prediction from movements (p = 0.0078, two-sided Wilcoxon sign rank test). In other words, the V1 activity evoked by sounds in individual trials followed a similar time course as the body movements observed in those trials.

Moreover, the behavioral model confirmed the intuition obtained from the correlations (Figure 4h): movements preceded the activity evoked by sounds in visual cortex. The kernel of a behavioral model fit to predict auditory PC1 during spontaneous activity showed that movement could best predicted neural activity occurring 25-50ms later (Suppl. Figure 12). This suggests that the activity evoked by sounds in visual cortex is driven by prior changes in internal and behavioral state.

## Discussion

These results confirm that sounds evoke activity in visual cortex^2-7^, but provide an alternative interpretation for this activity based on the widespread neural correlates of internal state and body movement^9,11-13,30^. We found that different sounds evoke different uninstructed body movements such as whisking, which reflect rapid changes in internal state. Crucially, we discovered that these movements are sufficient to explain the activity evoked by sounds in visual cortex.

Confirming this interpretation, we found that sound-evoked activity in visual cortex was independent of projections from auditory cortex. This result contrasts those of studies that ascribed the activity evoked by sounds in V1 to a direct input from auditory cortex. These studies used multiple methods: silencing of auditory cortex^2,5^; stimulation of its projections to visual cortex^2,3^; or transectomy of these projections^2^. However, the first two methods would interfere with auditory processing, and thus could affect sound-evoked behavior. We thus opted for transectomy^2^, which is less likely to modify behavior, and we performed bilateral recordings to have an internal control – the uncut side – within the same mice and with the same behavior. In contrast with the original study that introduced the transectomy^2^, but in accordance with our interpretation, these manipulations did not reduce sound-evoked activity in V1.

This difference in results with the previous study^2^ may be due to differences in methods. First, the previous study was conducted intracellularly and mostly in layers 2/3 (where sounds hyperpolarized cells, unlike in other layers where sounds increased spiking), whereas we recorded extracellularly in all layers (and observed mainly increases in spiking). Second, the previous study performed recordings hours after the transectomy, whereas we performed them days later. Third, the previous study anesthetized the mice, whereas we did not, a difference that can profoundly affect V1 activity^31^.

Our results indicate that sound-evoked activity is widespread in visual cortex and even in the hippocampal formation, and in both regions, it is low-dimensional. These properties echo those of movement-related activity, which is distributed all over the brain^12,13,21,30^ and low-dimensional^12^. We indeed found that movement-related activity even in the absence of sounds spanned essentially the same neural dimensions as sound-evoked activity. Moreover, the movements elicited by the sounds in each trial accurately predicted the subsequent sound-evoked activity. Taken together, these results suggest that, at least in our experiments, the sound-evoked activity had a behavioral origin.

This conclusion does not exclude the possibility of genuine auditory signals inherited from auditory cortex. After all, projections from auditory to visual cortex do exist, and they may perhaps carry auditory signals in other behavioral contexts, or in response to other types of stimuli. It is also possible that these projections represent a very sparse input, which influences only a minor fraction of V1 neurons and that we missed these neurons in our experiments.

However, our results show that distinguishing these putative auditory signals from the large contribution of internal state and behavior will require careful and systematic controls, which are rarely performed in passively listening mice. Some studies have controlled for eye movements^5^ or for overt behaviors such as licking^7^. However, previous studies may have overlooked the types of movements that we observed to correlate with neuronal activity, which were subtle twitches of the whiskers or the snout (see Suppl. Video 1). An exception is a study^19^ that explored the contribution of whisking to sound-evoked activity V1 neurons in layer 1 In agreement with our results, this study found that whisking explains a significant fraction of those neurons’ sound-evoked activity.

Our results do not imply that cortical activity is directly due to body movements; instead, cortical activity and body movements may both arise from changes in internal state. Consistent with this view, we found that sound-evoked activity in V1 was is low-dimensional, in sharp contrast to the highdimensional representation of visual stimuli^12^. This interpretation would explain some of the sound-evoked activity in visual cortex under anesthesia^2,3^, where movements are not possible, but state changes are common and difficult to control and monitor^32,33^.

Finally, these observations suggest that changes in states or behavior may also explain other aspects of neural activity that have been previously interpreted as being multisensory^8^. Stereotyped body movements can be elicited not only by sounds^15–18^ but also by images^34–38^ and odors^35,39^. These movements may be even more likely in response to natural stimuli ^18^ which are increasingly common in the field. Given the extensive correlates of body movement observed throughout the brain^12,13,20,30,40^ these observations reinforce the importance of monitoring behavioral state and body movement when interpreting sensory-evoked activity, in all brain areas and across contexts.

## Acknowledgements

We thank Philip Coen and Anwar Nunez-Elizalde for useful conversations and comments on the manuscript, Andrew Peters and Rebecca Terry for help with the widefield calcium imaging, Laura Funnell and Anne Ritoux for help with perfusion, Magdalena Robacha and Mickael Krumin for help with Brainsaw microscopy, and Yoh Isogai and Daniel Regester for providing the explantable methods. We also thank Giuliano Iurilli for advice on transectomies and for helpful discussions. This work was supported by the Wellcome Trust (grant 205093 to MC and KDH), by EMBO (ALTF 740-2019 fellowship to CB), and by the Sainsbury Wellcome Centre PhD program (TS and AL). MC holds the GlaxoSmithKline/Fight for Sight Chair in Visual Neuroscience.

## Author contributions

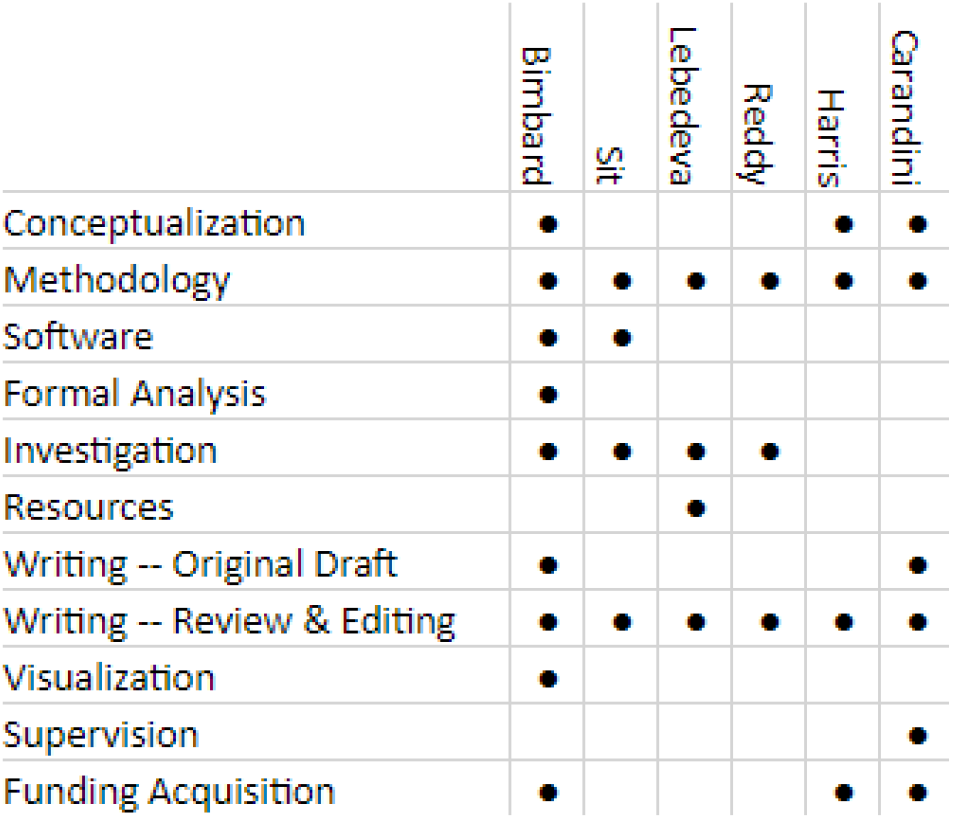

## Methods

Experimental procedures at UCL were conducted according to the UK Animals Scientific Procedures Act (1986) and under personal and project licenses released by the Home Office following appropriate ethics review.

### Surgery and recordings

Recordings were performed on 8 mice (6 male and 2 female), between 16 and 38 weeks of age. Mice were first implanted with a headplate designed for head-fixation under isoflurane anesthesia (1–3% in O_2_). After recovery, neural activity was recorded using Neuropixels 1.0 (n = 5) and 2.0 (n = 3, among which 2 had 4 shanks) probes implanted in left primary visual cortex (2.5 mm lateral, 3.5 mm posterior from Bregma, one probe per animal) and in the underlying hippocampal formation. In 5 of the mice the probes were implanted permanently or with a recoverable implant as described in Refs. ^23,41^ and in the remaining 3 they were implanted with a recoverable implant of a different design (Yoh Isogai and Daniel Regester, personal communication). Results were not affected by the implantation strategy. Sessions were automatically spike-sorted using Kilosort2 (www.github.com/MouseLand/Kilosort2^42^) and manually curated to select isolated single cells. The final number of cells was 611 in primary visual cortex (8 mice, 69 / 53 / 50 / 44 / 31 / 33 / 119 / 212 for each recording) and 230 in hippocampal formation (5 mice, 49 / 11 / 28 / 64 / 77 for each recording, mainly from dorsal subiculum and prosubiculum). Probe location was checked post-hoc by aligning it to the Allen Mouse Brain Atlas^43^ visually or through custom software (www.github.com/petersaj/AP_histology).

### Transectomy experiments

In 3 additional mice (all male, of 10, 21 and 22 weeks of age) we performed transectomies to cut the fibers running from auditory to visual cortex and followed them with bilateral recordings in visual cortex. Mice expressed GCamp6s in excitatory neurons (mouse 1 & 3: Rorb.Camk2tTA.Ai96G6s_L_001; mouse 2: tet0-G6s x CaMK-tTA) to enable us to monitor the normal activity of an intact visual cortex through widefield imaging (data not shown). Prior to headplate implantation, we used a dental drill (13,000 rpm) to perform a narrow rectangular (0.3 mm wide) craniotomy along the antero-posterior axis (from 1.6 mm posterior to 4.3 mm posterior) centered at 4.3 mm lateral to Bregma. To make the transectomy we then used an angled micro knife (angled 15°, 10315-12 from Fine Science Tools), mounted on a Leica digital stereotaxic manipulator with fine drive. Ensuring the skull was in a horizontal position (the difference between both DV coordinates did not exceed 0.1 mm), the knife was tilted 40° relative to the brain. The knife was inserted to a depth of 1.7 mm at the posterior end of the craniotomy, and slowly moved to the anterior end with the manipulator control. Any bleeding was stemmed by applying gelfoam soaked in cortex buffer. To protect the brain, we then applied a layer of Kwik-Sil (World Precision Instruments, Inc.) followed by a generous layer of optical adhesive (NOA 81, Norland Products Inc). Following this, a headplate was attached to the skull as described above and any exposed parts of the skull were covered with more optical adhesive. After a rest period of 1 week for recovery, we imaged the visual cortex under a widefield scope to confirm that it was healthy and responding normally to visual stimuli. Bilateral craniotomies were performed between 7 to 14 days following the transectomy, and acute bilateral recordings were acquired using 4-shank Neuropixels 2.0 probes targeting visual cortex over multiple days (3, 1 and 2 consecutive days in the three mice). The total number of cells was 1059 (ipsi) and 914 (contra) (per recording, ipsi;contra: 164;185 / 216;106 / 254;324 / 58;59 / 218;125 / 149;115). We imaged the brains using serial section^44^ two-photon^45^ microscopy. Our microscope was controlled by ScanImage Basic (Vidrio Technologies, USA) using BakingTray (https://github.com/SainsburyWellcomeCentre/BakingTray, https://doi.org/10.5281/zenodo.3631609). Images were assembled using StitchIt (https://github.com/SainsburyWellcomeCentre/StitchIt, https://zenodo.org/badge/latestdoi/57851444). Probe location was checked using brainreg^26–28^, showing that most recordings were in area V1, and partially VISpm and VISl. The exact location of the probe in visual cortex did not affect the results so we pooled all areas together under the name of VIS.

### Stimuli

In each session, mice were presented with a sequence of audio, visual or audiovisual movies. The stimuli consisted of all combinations of auditory and visual streams extracted from a set of 11 naturalistic movies depicting the movement of animals such as cats, donkeys and seals, from the AudioSet database^24^. An additional visual stream consisted of a static full-field gray image and an additional auditory stream contained no sound. Movies lasted for 4 s, and were separated by an inter-trial interval of 2 s. The same randomized sequence of movies was repeated 4 times during each experiment, with the second and third repeat separated by a 5 min interval.

The movies were gray-scaled, spatially re-scaled to match the dimensions of a single screen of the display, and duplicated across the three screens. The visual stream was sampled at 30 frames per second. Visual stimuli were presented through three displays (Adafruit, LP097QX1) each with a resolution of 1024 by 768 pixels. The screens covered approximately 270 x 70 degrees of visual angle, with 0 degree being directly in front of the mouse. The screens had a refresh rate of 60 frames per second and were fitted with Fresnel lenses (Wuxi Bohai Optics, BHPA220-2-5) to ensure approximately equal luminance across viewing angles.

Sounds were presented through a pair of Logitech Z313 speakers placed below the screens. The auditory stream was sampled at 44.1 kHz with 2 channels and was scaled to a sound level of −20 decibels relative to full scale.

*In situ* sound intensity and spectral content was estimated using a calibrated microphone (GRAS 40BF 1/4” Ext. Polarized Free-field Microphone) positioned where the mice sit, and reference loudness was estimated using an acoustic calibrator (SV 30A, Suppl. Figure 1). Mice were systematically habituated to the rig through 3 days of familiarizing with the rig’s environment and head-fixation sessions of progressive duration (from 10 min to an hour). They were not habituated to the specific stimuli before the experiment. Two exceptions were the transectomy experiments, where mice were presented with the same protocol across the consecutive days of recordings (so a recording on day 2 would mean the mouse had been through the protocol one already), and in specific control experiments not shown here (n = 2 mice). Presentation of the sounds over days (from 2 to 5 days) did not alter the observed behavioral and neural responses (n = 2 transectomy mice + 2 control mice).

### Videography

Eye and body movements were monitored by illuminating the subject with infrared light (830 nm, Mightex SLS-0208-A). The right eye was monitored with a camera (The Imaging Source, DMK 23U618) fitted with zoom lens (Thorlabs MVL7000) and long-pass filter (Thorlabs FEL0750), recording at 100 Hz. Body movements (face, ears, front paws, and part of the back) were monitored with another camera (same model but with a different lens, Thorlabs MVL16M23) situated above the central screen, recording at 40 Hz for the experiments in V1 and HPF (Figure 1 & Figure 2) and 60Hz for the transectomy experiments (Figure 3). Video and stimulus time were aligned using the strobe pulses generated by the cameras, recorded alongside the output of a screen-monitoring photodiode and the input to the speakers, all sampled at 2,500 Hz. To compute the Singular Value Decompositions of the face movie and to fit pupil area and position, we used the *facemap* algorithm^12^ (www.github.com/MouseLand/facemap).

### Behavior-only experiments

In order to test for the influence of basic acoustic properties on movements, we ran behavior-only experiments (i.e., only with cameras filming the mice, and no electrophysiology, Suppl. Figure 5) on 8 mice in which we played i) white noise of various intensities; ii) pure tones of various frequencies; iii) white noise coming from various locations. In contrast with the previous experiments, auditory stimuli were presented using an array of 7 speakers (102-1299-ND, Digikey), arranged below the screens at 30° azimuthal intervals from −60° to +60° (where −90°/+90° is directly to the left/right of the subject). Speakers were driven with an internal sound card (STRIX SOAR, ASUS) and custom 7-channel amplifier (http://maxhunter.me/portfolio/7champ/). As in the previous experiments, *in situ* sound intensity and spectral content was estimated using a calibrated microphone (GRAS 40BF 1/4” Ext. Polarized Free-field Microphone) positioned where the mice sit, and reference loudness was estimated using an acoustic calibrator (SV 30A). Body movements were monitored with a Chameleon3 camera (CM3-U3-13Y3C-S-BD, Teledyne FLIR) recording at 60Hz. The movie was then processed with *facemap*.

The effect of each factor was then quantified using repeated-measures ANOVA with either the sound loudness, frequency, or location as a factor.

### Data processing

For each experiment, the neural responses constitute a 5-dimensional array **D** of size *N_t_* time bins x *N_v_* videos x *N_a_* sounds x *N_r_* repeats x *N_c_* cells. The elements of this matrix are the responses *D_tvarc_* measured at time *t*, in video *v*, sound *a*, repeat *r*, and cell *c*. **D** contains the binned firing rates (30 ms bin size) around the stimulus onset (from 1 s before onset to 3.8 s after onset), smoothed with a causal half gaussian filter (standard deviation of 43 ms), and z-scored for each neuron.

Pupil area and eye position were baseline-corrected to remove the slow fluctuations and focus on the fast, stimulus-evoked and trial-based fluctuations: the mean value of the pupil area or eye position over the second preceding stimulus onset was subtracted from each trial. Signed eye motion (horizontal and vertical) was computed as the difference of the eye position between time bins. The unsigned motion was obtained as the absolute value of the signed motion. The global eye motion was estimated as the absolute value of the movement in any direction (L2 norm). Eye variables values during identified blinks were interpolated based on their values before and after the identified blink. Body motion variables were defined as the first 128 body motion PCs. Both eye-related and body-related variables were then binned similarly to the neural data. We note that the timing precision for the face motion is limited by both the camera acquisition frame rate (40 fps, not aligned to stimulus onset), and the binning used here (30 ms bins, aligned on stimulus onset). Thus, real timings can differ by up to 25 ms.

All analyses that needed cross-validation (test-retest component covariance, decoding, prediction) were performed using a training set consisting of half of the trials (odd trials) and a test set based on the other half (even trials). Models were computed on the train set and tested on the test set. Then test and train sets were swapped, and quantities of interest were averaged over the two folds.

To estimate the correlation of the sound-evoked time courses across mice, the variable of interest was split between training and test set, averaged over all trials (e.g., for sound-related activity, over videos and repeats), and the Pearson correlation coefficient was computed between the training set activity for each mouse and the test set activity of all mice (thus giving a cross-validated estimate of the auto- and the cross-correlation). Averages were obtained by Fisher’s Z-transforming each coefficient, averaging, and back-transforming this average.

### Marginalization

To isolate the contribution of videos or sounds in the neural activity we used a marginalization procedure similar to the one used in factorial ANOVA. By *D_tvarc_* we denote the firing rate of cell *c* to repeat *r* of the combination of auditory stimulus *a* and visual stimulus *v*, at time *t* after stimulus onset. The marginalization procedure decomposes *D_tvarc_* into components that are equal across stimuli, related to videos only, related to sounds only, related to audiovisual interactions, and noise:

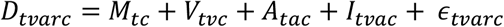

The first term is the mean of the population activity across videos, sounds, and repeats:

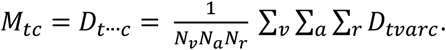

where dots in the second term indicate averages over the missing subscripts, and *N_v_, N_a_, N_r_* denote the total number of visual stimuli, auditory stimuli, and repeats.

The second term, the video-related component, is the average of the population responses over sounds and repeats, relative to this mean response:

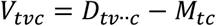

Similarly, the sound-related component is the average over videos and repeats, relative to the mean response:

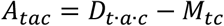

The audiovisual interaction component is the variation in population responses that is specific to each pair of sound and video:

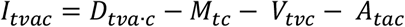

Finally, the noise component is the variation across trials:

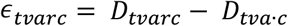

In matrix notation, we will call **A, V**, and **I** the arrays with elements *A_tac_, V_tvc_*, and *I_tvac_* and size *N_t_* × *N_a_* × *N_c_, N_t_* × *N_v_* × *N_c_* and *N_t_* × *N_v_* × *N_a_* × *N_c_*.

### Dimensionality reduction

The arrays of sound-related activity **A**, of video-related activity **V**, and of audiovisual interactions **I**, describe the activity of many neurons. To summarize this activity, we used cross-validated Principal Component Analysis^14^ (cvPCA). In this approach, principal component projections are found from one half of the data, and an unbiased estimate of the reliable signal variance is found by computing their covariance with the same projections on a second half of the data.

We illustrate this procedure on the sound-related activity. In what follows, all arrays, array elements, and averages (e.g. **A**, *A_tac_, A_t.c_*) refer to training-set data (odd-numbered repeats), unless explicitly indicated with the subscript *test* (e.g. ***A**_test_, A_tac;test_, A_t.c.test_*).

We first isolate the sound-related activity **A** as described above from training set data (odd-numbered trials). We reshape this array to have two dimensions *N_t_N_a_* × *N_c_*; and perform PCA:

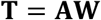

where **T** (*N_t_N_a_* × *N_p_*) is a set of time courses of the top *N_p_* principal components of **A**, and **W** is the PCA weight matrix (*N_c_* × *N_p_*).

For cvPCA analysis, we took *N_p_* = *N_c_* to estimate the amount of reliable stimulus-triggered variance in each dimension (Fig. 2f,i; Supp. Fig. 2). We computed the projections of the mean response over a test set of even-numbered trials, using the same weight matrix: **T**_*test*_ = **A**_*test*_**W** and evaluated their covariance with the training-set projections:

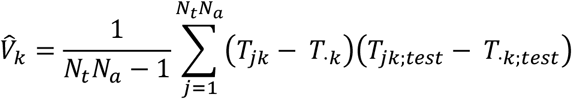

This method provides an unbiased estimate of the stimulus-related variance of each component^14^. Analogous methods were used to obtain the signal variance for principal components of the visual response and interaction, by replacing **A** with **V** or **I** (Supp. Fig 2). The cvPCA variances were normalized either by the sum for all auditory dimensions (e.g., Figure 2h,j), or the sum for all dimensions from video-related, sound-related and interaction-related decompositions (Suppl. Figure 2).

To determine if a cvPCA dimension had variance significantly above 0, we used a shuffling method. The shuffling was done by changing the labels of both the videos and the sounds for each repeat. We performed this randomization 1,000 times and chose a component to be significant if its test-retest covariance value was above the 99^th^ percentile of the shuffled distribution. We defined the dimensionality as the number of significant components. For the video-related activity, we found an average of 76 significant components (± 22, s.e., n = 8 mice). As expected, this number grew with the number of recorded neurons^14^ (data not shown). For the sound-related activity, instead, we found only 4 significant components on average (± 0.9, s.e., n = 8 mice). For the interactions between videos and sounds, finally, we found zero significant components (0 ± 0, s.e., n = 8 mice) indicating that the population responses did not reflect significant interactions between videos and sounds.

For visualization of PC time courses (Figure 1, Figure 2 & Figure 3), the weight matrices **W** were computed from the training set but the projection of the full dataset was used to compute the time courses of the first component.

### Decoding

Single-trial decoding for video- or sound-identity was performed using a template-matching decoder applied to neural or behavioral data. In this description, we will focus on decoding sound identity from neural data. The data were again split into training and test sets consisting of odd and even trials. Both test and trained trials contained a balanced number of trials for each sound.

When decoding sound-related neural activity (Figure 1, Figure 2, and Figure 3), we took *N_p_* = 4, so the matrix **T** containing PC projections of the mean training-set sound-related activity had size *N_t_N_a_* × 4; using more components did not affect the results. To decode the auditory stimulus presented on a given test-set trial, we first removed the video-related component by subtracting the mean response to the video presented on that trial (averaged over all training-set trials) We then projected this using the training-set weight matrix **W** to obtain a *N_t_* × 4 timecourse for the top auditory PCs, and found the best-matching auditory stimulus by comparing to the mean training-set timecourses for each auditory stimulus using Euclidean distance. A similar analysis was used to decode visual stimuli, using *N_p_ =* 30 components in visual cortex and *N_p_* = 4 in the hippocampal formation.

To decode the sound identity from behavioral data, we used the z-scored eye variables (pupil area and eye motion in Suppl. Figure 6), or the first 128 principal components of the motion energy of the face movie (Figure 4) and performed the template-matching the same way as the with the neural data.

The significance of the decoding accuracy (compared to chance) was computed by performing a right-sided Wilcoxon sign rank test to compare to chance level (1/12), treating each mouse as independent. The comparison between video identity and sound identity decoding accuracy was computed by performing a paired two-sided Wilcoxon sign rank test across mice.

### Encoding

To predict neural activity from stimuli/behavioral variables (“encoding model”; Figure 4, Suppl. Figure 8), we again started by extracting audio- or video-related components and performing Principal Component Analysis, as described above, however this time the weight matrices were computed from the full dataset rather than only the training set. Again, we illustrate by describing how sound-related activity was predicted, for which we kept *N_p_* = 4 components; video-related activity was predicted similarly but with *N_p_* = 30 in visual cortex and *N_p_* = 4 in the hippocampal formation.

We predicted neural activity using linear regression. The target **Y** contained the marginalized, sound-related activity on each trial, projected onto the top 4 auditory components: specifically, we compute *D_tvarc_* – *M_tc_* – *V_tvc_* reshape to a matrix of size *N_t_N_v_N_a_N_r_* × *N_c_*, and multiply by the matrix of PC weights **W**. We predicted **Y** by regression: **Y** ≈ **XB**, where **X** is a feature matrix and **B** are weights fit by cross-validated ridge regression.

The feature matrix depended on the model. To predict from sensory stimulus identity (see ‘Auditory predictors’ in Suppl. Figure 8), **X** had one column for each combination of auditory stimulus and peristimulus timepoint, making *N_a_N_t_* =1,524 columns, *N_t_N_v_N_a_N_r_* rows, and contained 1 during stimulus presentations in a column reflecting the stimulus identity and peristimulus time. With this feature matrix, the weights **B** represent the mean activity time course for each dimension and stimulus, and estimation is equivalent to averaging across the repeats of the train set. It is thus equivalent to a test-retest estimation and is not a model based on acoustic features of the sounds.

To predict from behavior, we used features for pupil area, pupil position (horizontal and vertical), eye motion (horizontal and vertical -- signed and unsigned), global eye motion (L2 norm of x and y motion, unsigned), blinks (thus 9 eye-related predictors) and the first 128 face motion PCs, with lags from −100 ms to 200 ms (thus 12 lags per predictor, 1,644 predictors total, see ‘Eye predictors’ and ‘Body motion predictors’ in Suppl. Figure 8). As for the neural activity target matrix **Y**, all behavioral variables were first marginalized to extract the sound-related modulations. To predict from both stimulus identity and behavior, we concatenated the feature matrices, obtaining a matrix with 3,168 columns. The beginning and end of the time course for each trial were padded with NaNs (12 - the number of lags - at the beginning and end of each trial, to avoid cross-trial predictions by temporal filters. Thus, the feature matrix has (*N_t_* + 24)*N_v_N_a_N_r_* rows. A model with the eye variables only, and a model with the face motion variables only was also constructed (Suppl. Figure 9). Note that in the case of mice for which the eye wasn’t recorded (2 out of the 8 mice, and all transectomy experiments), the behavioral model contained only the body motion variables.

We used ridge regression to predict the single trial version of **Y** from **X** on the training set. The best lambda parameter was selected using a 3-fold cross-validation within the training set.

To measure the accuracy of predicting trial-averaged sound-related activity (Figure 4), we averaged the *N_t_N_v_N_a_N_r_* × *N_p_* activity matrix **Y**_*test*_ over all test-set trials of a given auditory stimulus, to obtain a matrix of size *N_t_N_a_* × *N_c_*, and did the same for the prediction matrix **X**_*test*_**B**, and evaluated prediction quality by the elementwise Pearson correlation of these two matrices.

To evaluate predictions of trial-to-trial fluctuations (Suppl. Figure 11b,c), we computed a “noise” matrix of size *N_t_N_v_N_a_N_r_* × *N_p_* by subtracting the mean response to each sound: *Y_tvarp;test_* – *Y_t·a·p; test_*, performed the same subtraction on the prediction matrix **X**_*test*_**B**, and evaluated prediction quality by the elementwise Pearson correlation of these two matrices. Again, the average was obtained by Fisher’s Z-transforming each coefficient, averaging, and back-transforming this average.

To visualize the facial areas important to explain neural activity (Suppl. Figure 9), we reconstructed the weights of the auditory PC1 prediction in pixel space. Let 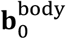 (1 × 128) be the weights predicting neural auditory PC1 at lag 0 from each of the 128 body motion PCs. Let **ω** (128 × total number of pixels in the video) be the weights of each of these 128 face motion PCs in pixel space (as an output of the *facemap* algorithm). We obtained an image **I** of the pixel-to-neural weights by computing 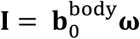.

Finally, to explore the timing relationship between movement and neural activity, we looked at the cross-correlogram of the motion PC1 and the auditory PC1 during the spontaneous (no stimulus) period (Figure 4, Suppl. Figure 7). The auditory PC1 was found by computing its weights without cross-validation. To maximize the temporal resolution, the regression analysis was performed on the spikes sampled at the rate of the camera acquisition (40 fps, thus 25ms precision). We then computed the lag associated in the cross-correlogram, which showed that movement preceded neural activity by 25-50ms. To avoid errors induced by “large” cross-correlograms due to autocorrelation of the two signals, we also performed a ridge regression of the auditory PC1 from the motion PCs during the spontaneous period and looked at the peak of the weights of motion PC1 to predict auditory PC1 (Suppl. Figure 12).

### Movement- and sound-related subspaces overlap

To quantify the overlap between the movement- and the sound-related subspaces of neural activity in V1, we computed how much of the sound-related variance the movement-related subspace could explain^12^. We first computed the movement-related subspace by computing a reduced-rank regression model to predict the neural activity matrix **S** (*T* × *N_c_*, with *T* being the number of time points) from the motion components matrix **M** with lags (*T* × 128*21=2,688 lags) during the spontaneous period (no stimulus), both binned at the face video frame rate (40 or 60Hz). This yields a weight matrix **B** (2,688 × *N_c_*) so that: **S** ≈ **MB**. The weight matrix **B** factorizes as a product of two matrices of sizes 2,688 × *r* and *r* × *N_c_*, with *r* being the rank of the reduced-rank regression. The second part of this factorization, the matrix of size *r* × *N_c_* of which transpose we call **C** (*N_c_* × *r*), forms an orthonormal basis of the movement-related subspace of dimensionality *r*. Here, we chose *r* = 4 to match the size of the sound related subspace, but the results were not affected by small changes in this value. Then, we projected the sound-related activity of the train set **A** and the test set **A**_*test*_ onto **C** and measured the covariance of these projections for each dimension of the movement-related subspace. This is similar to the cvPCA performed above to find the variance explained by auditory PCs, except the components are here the ones most-explained by behavior, and not by sound. The overlap between the movement-related and the sound-related subspaces was finally quantified as the ratio of the sound-related variance explained by the first 4 components of each subspace.

We note that the fact that the overlap between the sound-related subspace and the behavior-related subspace isn’t 100% may come from the noise in estimating the behavior-related subspace, which relies on the spontaneous period only which was less than 25 min.

### Transectomy quantification

To visualize and estimate the extent of the transectomy, we used the software *brainreg*^26-28^ to register the brain to the Allen Mouse Brain Reference Atlas^43^, and manually trace the contours of the cut using brainreg-segment. The cut was identified visually by observing the massive neuronal loss (made obvious by a loss of fluorescence) and scars.

To estimate the extent of the fibers that were cut by the transectomy, we took advantage of the large-scale connectivity database of experiments performed by the Allen Brain Institute (Allen Mouse Brain Connectivity Atlas^25^). Using custom Python scripts, we selected and downloaded the 53 experiments where injections were performed in the auditory cortex and projections were observed in visual cortex (we subselected areas V1, VISpm and VISl as targets, since these were where the recordings were performed). We used the fiber tractography data to get the fibers’ coordinates in the reference space of the Allen Mouse Brain Atlas, to which was also aligned the actual brain and the cut reconstruction. Using custom software, we selected only the fibers of which terminal were inside or within 50 μm of either ipsilateral or contralateral visual cortex. We identified the cut fibers as all fibers that were passing inside or within 50 μm of the cut. Because auditory cortex on one side sends projections to both sides (yet much more to the ipsilateral side), cutting the fibers on one side could also affect responses on the other side. We thus quantified the auditory input to each visual cortex as the number of intact fibers coming from both auditory cortices, with one side being cut and the other being intact. We then made the hypothesis that the size of the responses, or more generally the variance explained by sounds in both populations, would linearly reflect these “auditory inputs”. We then compared the sound-related variance on the cut side to its prediction from the sound-related variance on the uncut side. This provided an internal control, with the same sounds and behavior. We took the sound-related variance as the cumulative sum of the variance explained by the first 4 auditory PCs, on both sides.

We then used *brainrender*^29^ to visualize all results.

## Code & data availability

All code and data will be available upon publication.

## Supplementary Figures

**Suppl. Figure 1.**
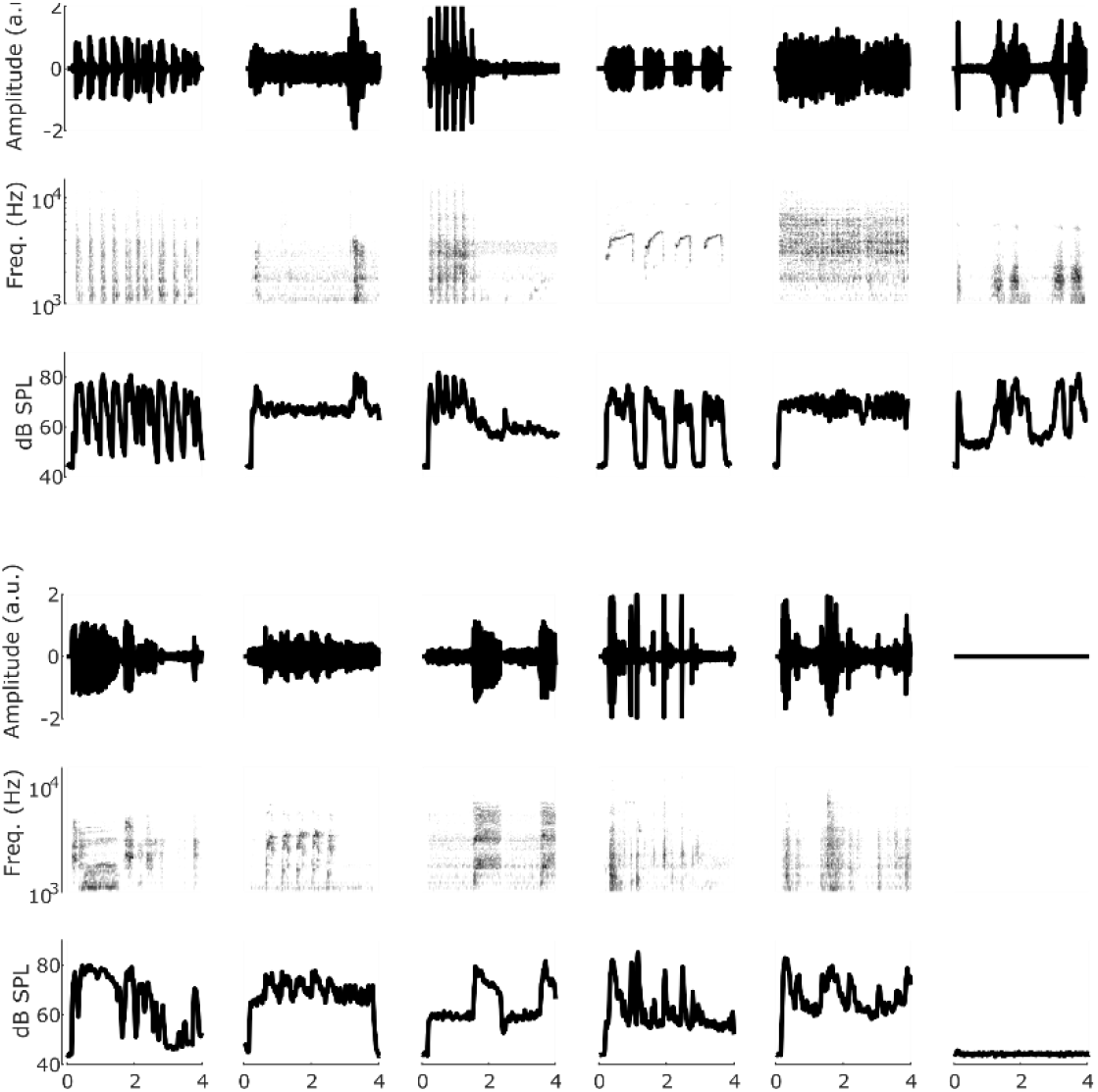
Naturalistic sounds used in this study: spectral content and loudness. For each sound is displayed: top: amplitude; middle: frequency spectrum; bottom: loudness.

**Suppl. Figure 2.**
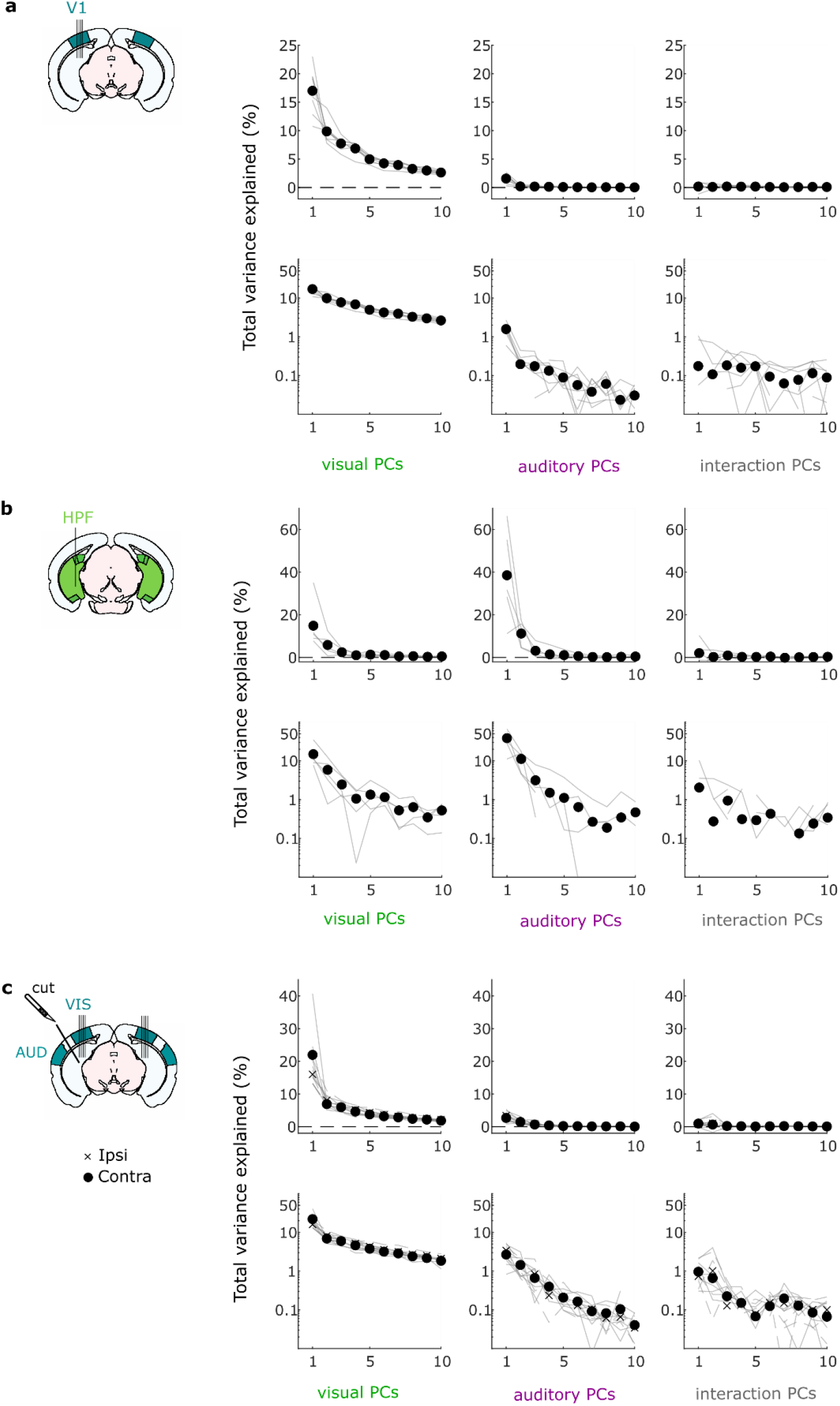
Neural responses are largely visually driven. **a.** Top: Total variance explained (normalized test-retest covariance) for videos PCs (*left*), auditory PCs (*middle*) and interactions PCs (*right*), for all 8 recordings in V1 (thin lines) and their average (filled dots). The test-retest covariance for each component was normalized by the total amount of test-retest covariance across components and video/sound/interaction conditions to show comparable proportions. Bottom: Same as top but with a logarithmic scale for the y-axis. Negative values are not displayed. **b.** Same as **a** but with the 5 recordings from the HPF. **c.** Same as **a** but with the 12 recordings from the visual cortices ipsilateral (6, crosses) and contralateral (6, filled dots) to the cut.

**Suppl. Figure 3.**
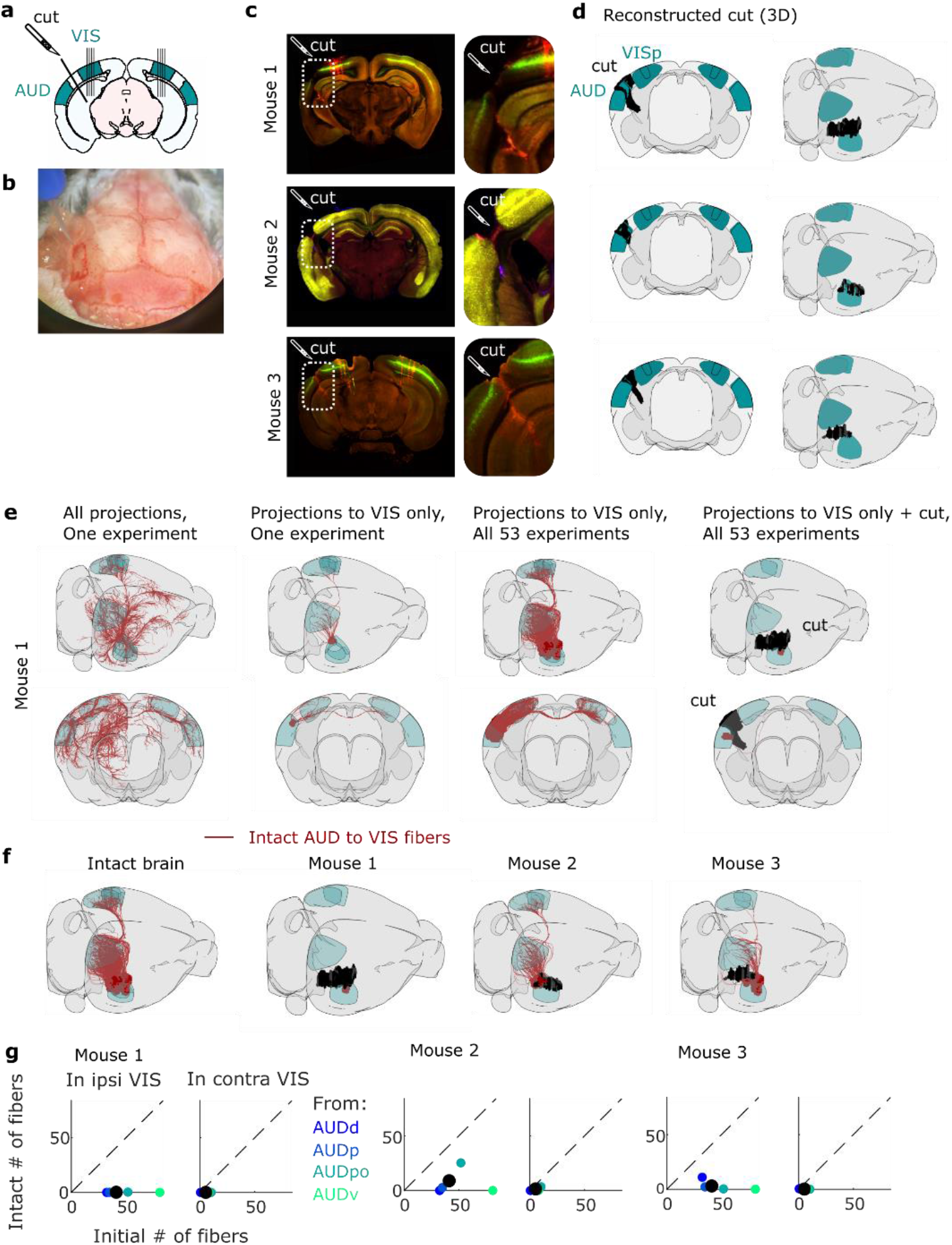
Transectomy experiments cut a large fraction of the auditory to visual cortex fibers. **a.** Schematics of the transectomy experiments: the connections between auditory (AUD) and visual (VIS) cortex are cut on one side. Simultaneous, bilateral recordings are performed in visual cortex, in both hemispheres. **b.** Picture from above of the mouse skull during surgery, with a craniotomy performed on the left side, and the fine scalpel used for craniotomies. **c.** Histology of the three mice’ brain, showing the cut (inset), and the recordings (DiI and DiO staining, mainly visible on mouse 1 and 3). **d.** 3D visualization of the reconstruction of the cut, shown from a coronal view (*left*) or from above/the side (right). **e.** Procedure to quantify of the extent of the cut. Far left: Fibers tracks from the auditory cortex to all brain regions in one experiment (taken from the Allen Mouse Brain Connectivity atlas^25^, see Methods); middle left: projections to the visual cortex only, from the same experiment; middle right: projections to the visual cortex only across all experiments; far right: projections to the visual cortex across all experiments, after the cut. Only the intact fibers are shown. **f**. Theoretically intact fibers for the 3 different mice. **g.** Quantification of the number of initial vs. intact fibers for each of the 53 experiments, grouped by injection sites, for each mouse, in both ipsi- and contralateral visual cortex (left and right plots). The color of the dot indicates where the injection was performed, as indicated by the Allen Mouse Brain Connectivity Atlas metadata. The black dot shows the average over all experiments.

**Suppl. Figure 4.**
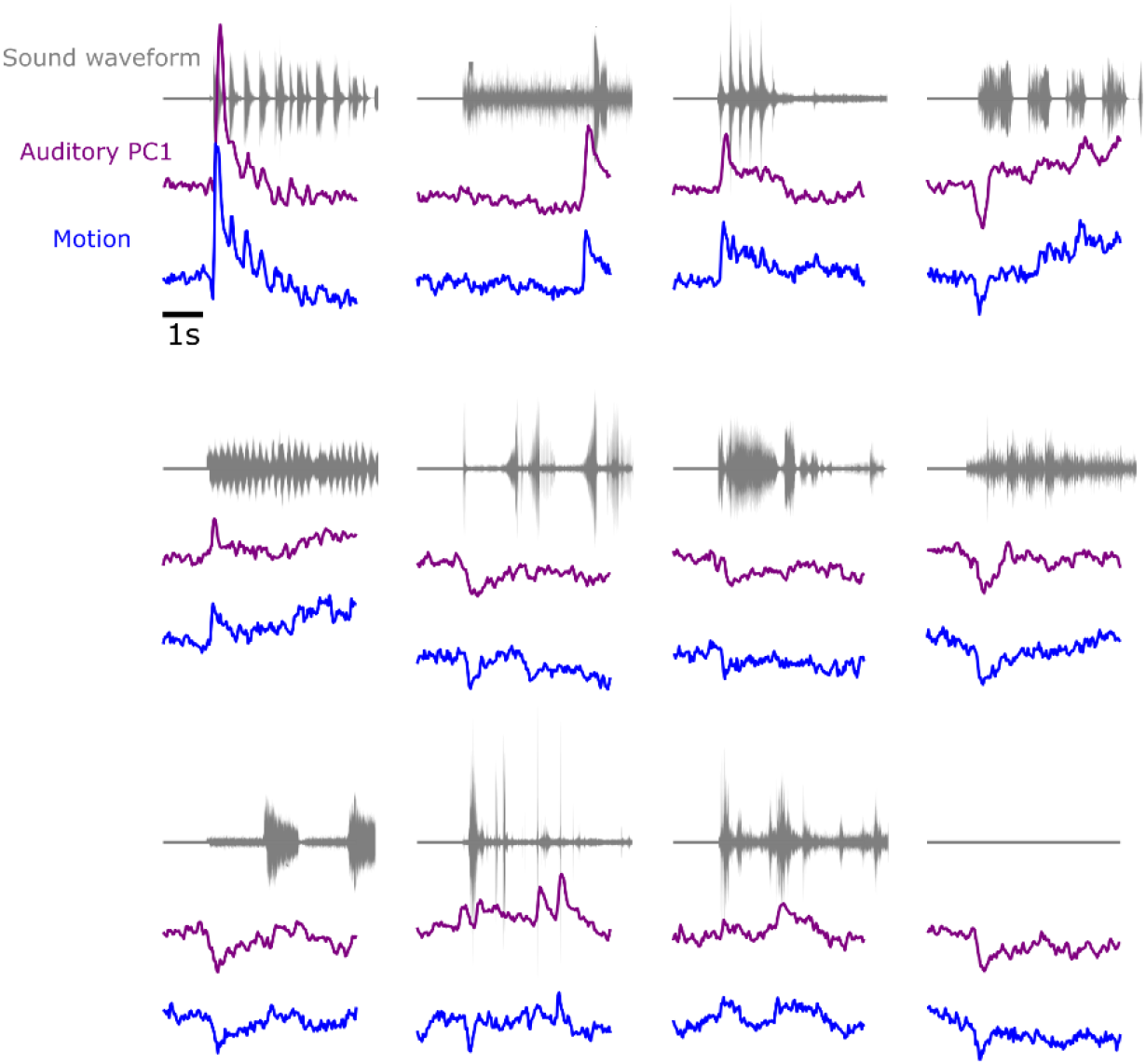
Neural and behavioral responses differ across sounds but resemble each other. Responses along neural auditory PC1 from V1 (purple), and motion energy (blue) sampled at 30 ms time bins for all sounds. Responses are averaged over trials, videos, and mice, and z-scored. The top trace (gray) shows the envelope of the corresponding sound. As in Figures 1, 2 and 3, these responses are expressed relative to the grand average over sounds and videos; this explains the negative deflections seen in the responses to the blank stimulus.

**Suppl. Figure 5.**
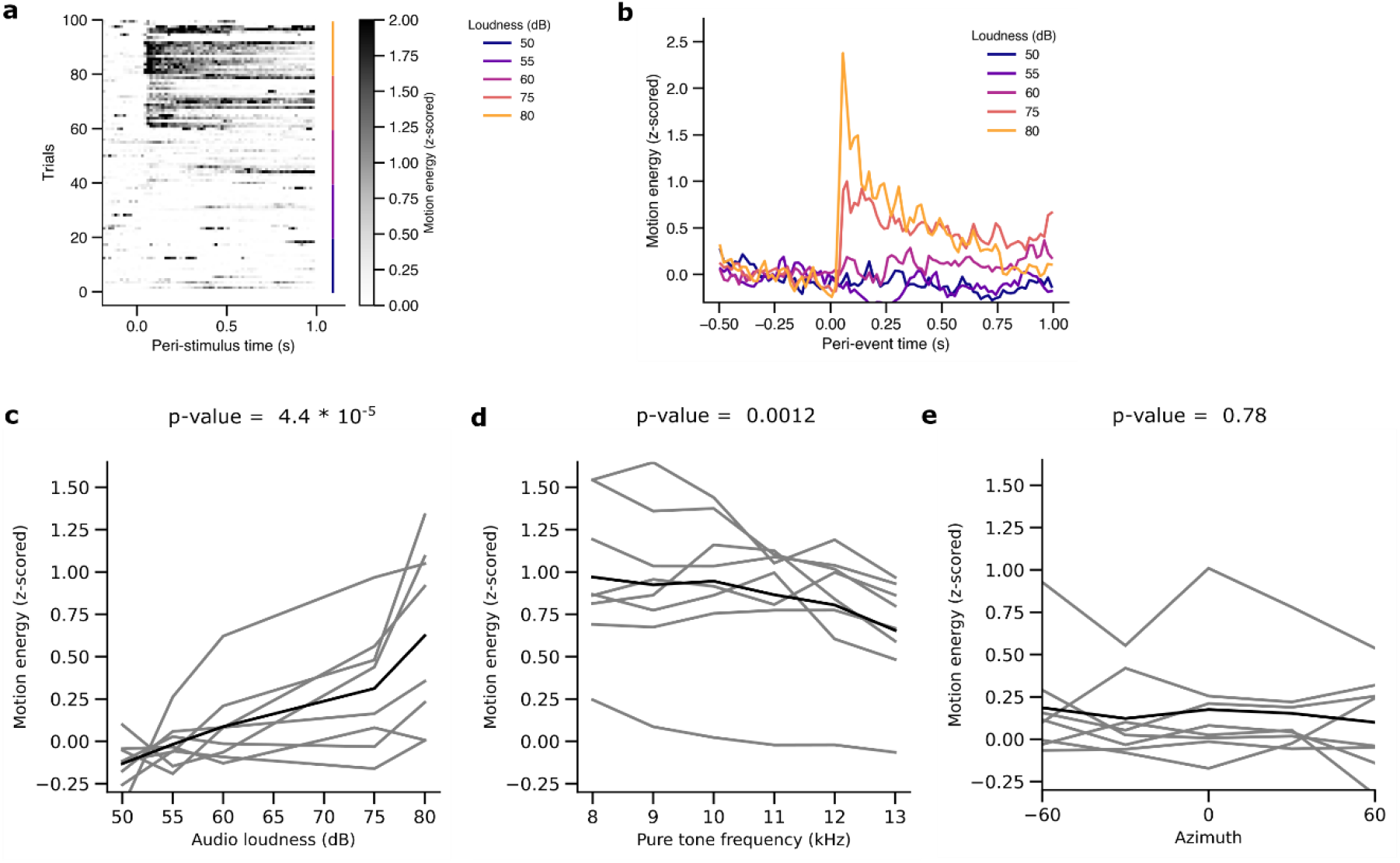
Loudness is the main driver of uninstructed behavioral responses. **a.** Raster of the average motion of an example mouse in response to white noise bursts of different loudness (from 50 to 80dB SPL). **b.** Peri-stimulus time histograms of the average motion for the same example mouse for different sound volumes. **c.** Average motion energy of 6 different mice as a function of loudness. **d**. Same as **c**, but for a pure tone of different frequencies (60dB). **e**. Same as **c**, but for a white noise burst played from different azimuthal locations (80dB).

**Suppl. Figure 6.**
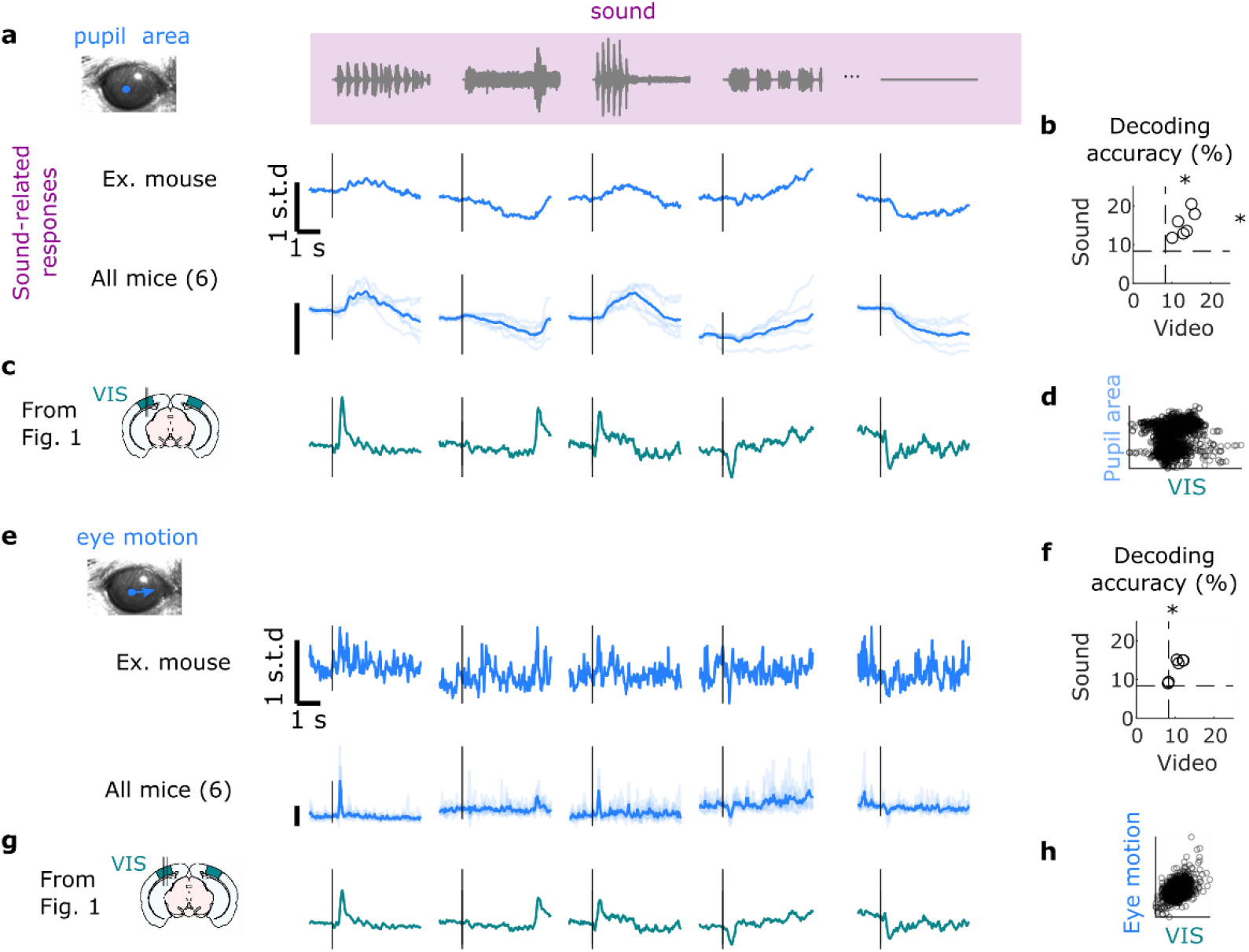
Sounds trigger changes in arousal and eye movements. **a.** Sound-related responses changes in arousal as measured by pupil diameter, for one example mice, and all mice (6 out of 8 were monitored with an eye camera). **b.** Decoding of video and sound identity using pupil area. **c.** Comparison with the timecourse of the neural auditory PC1 from Fig1. **d**. Scatter plot of the correlation. **f-h.** Same as **a-d.** but for eye movements.

**Suppl. Figure 7.**
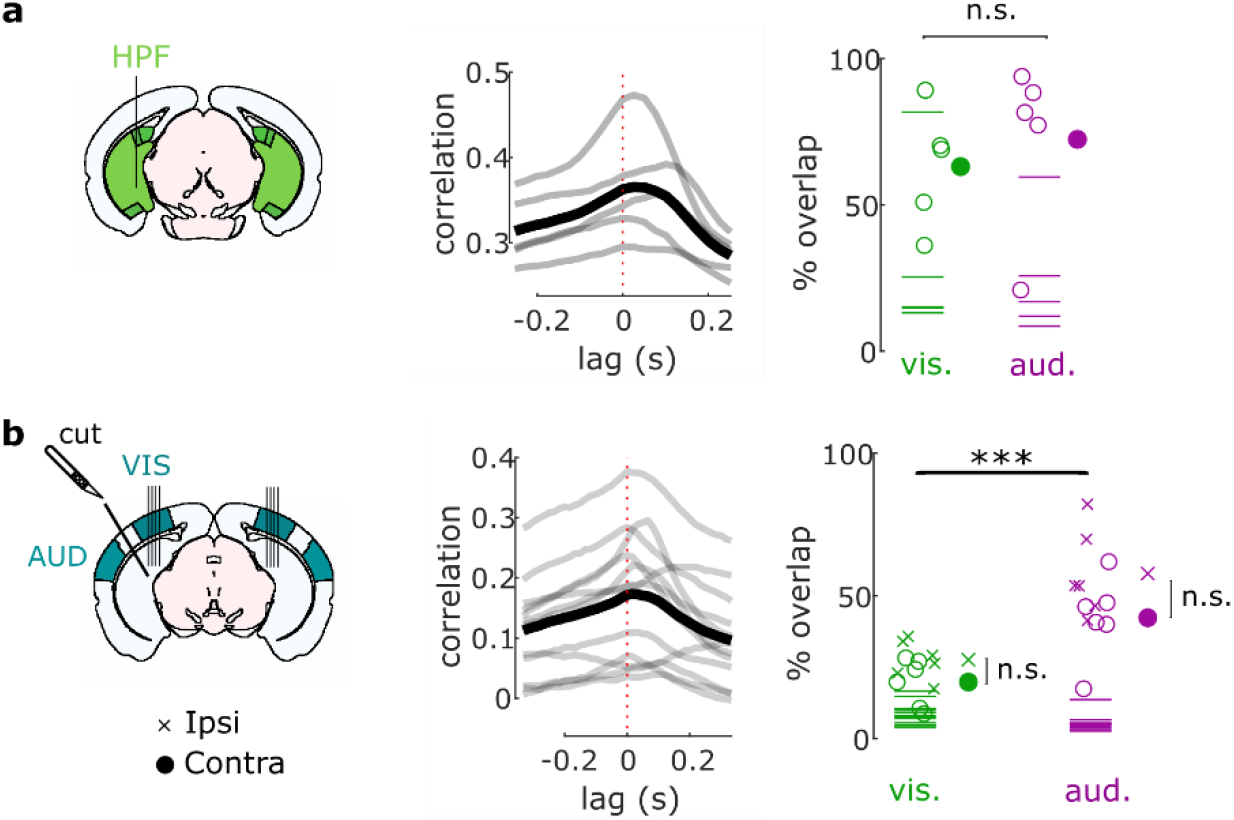
Movement preceded neural activity in the HPF and independently of auditory inputs to visual cortex, and behavior-related subspace overlapped with sound-related subspace. **a.** *Left*: Cross-correlogram of the motion energy and the neural activity on the auditory PC1 during the spontaneous period (grey lines, individual mice, thick black line, average). A positive lag means that movement precedes neural activity. *Right:* Percentage of overlap between the sound-related (or video-related) and the behavior-related subspace for each mouse (empty dots) and all mice (filled dot). Dashed lines show the significance threshold (95^th^ percentile of the overlap with random dimensions) for each mouse. **b**. Same as **a** for the recordings in visual cortex after a transectomy.

**Suppl. Figure 8.**
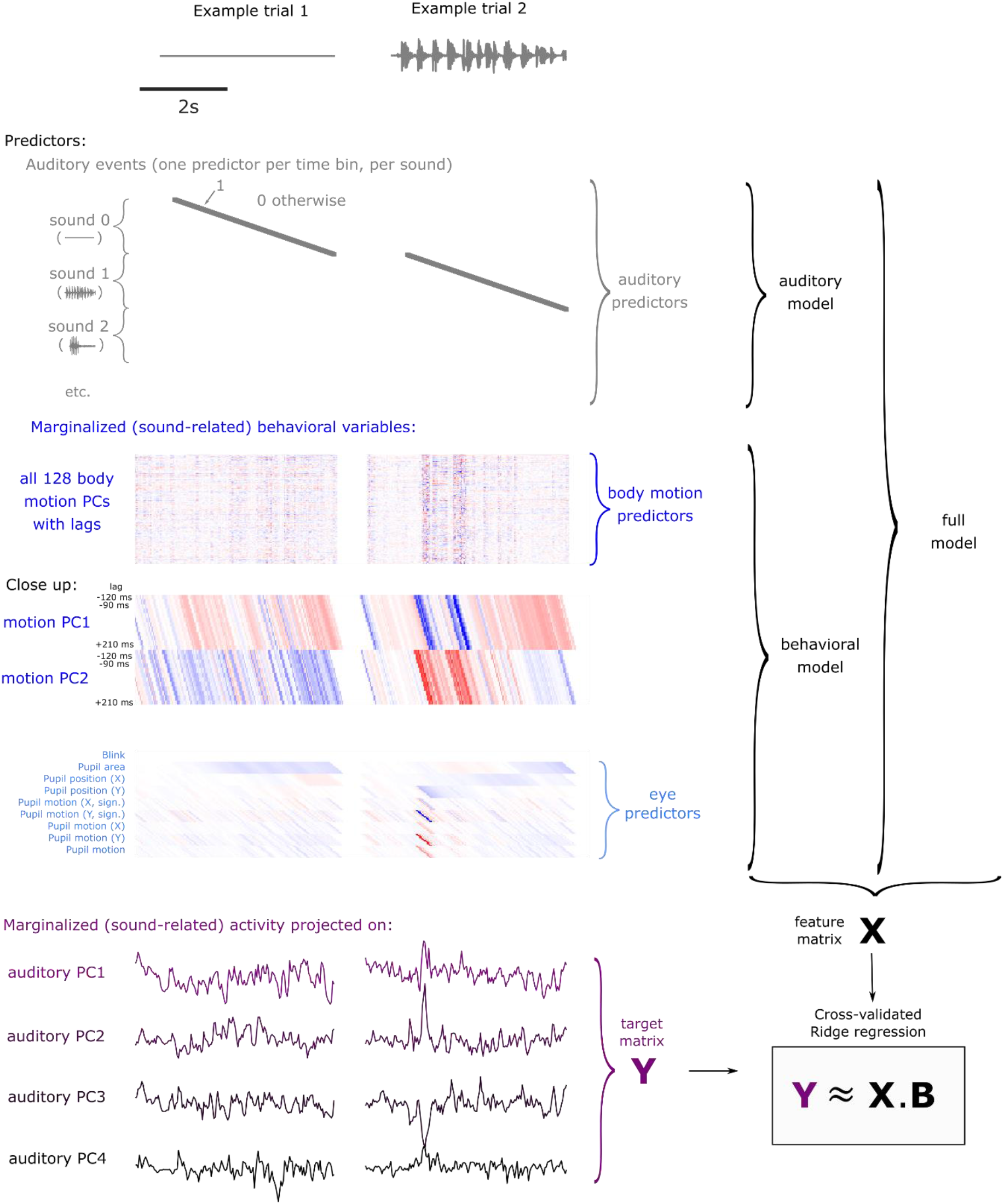
Structures of the models. The feature matrix **X** depends on the model. In the auditory model, there are as many predictors as the number of sounds multiplied by the number of time bins during which the sound is played (grey dots show where the value is 1, and 0 otherwise). In the behavioral model, the predictors consist of all 128 motions PCs, with various eye variables (see Methods), with 12 different lags (from −120 to +210ms). A close up for the first 2 motion PCs allows for a better visualization of the predictors with different lags. The target matrix **Y** is composed of the projections of the marginalized activity (sound responses only) onto the sound-related subspace (first 4 auditory PCs). Finally, the full model combines both the auditory and the behavioral predictors. The model is then fitted using 3-fold cross-validated ridge regression.

**Suppl. Figure 9.**
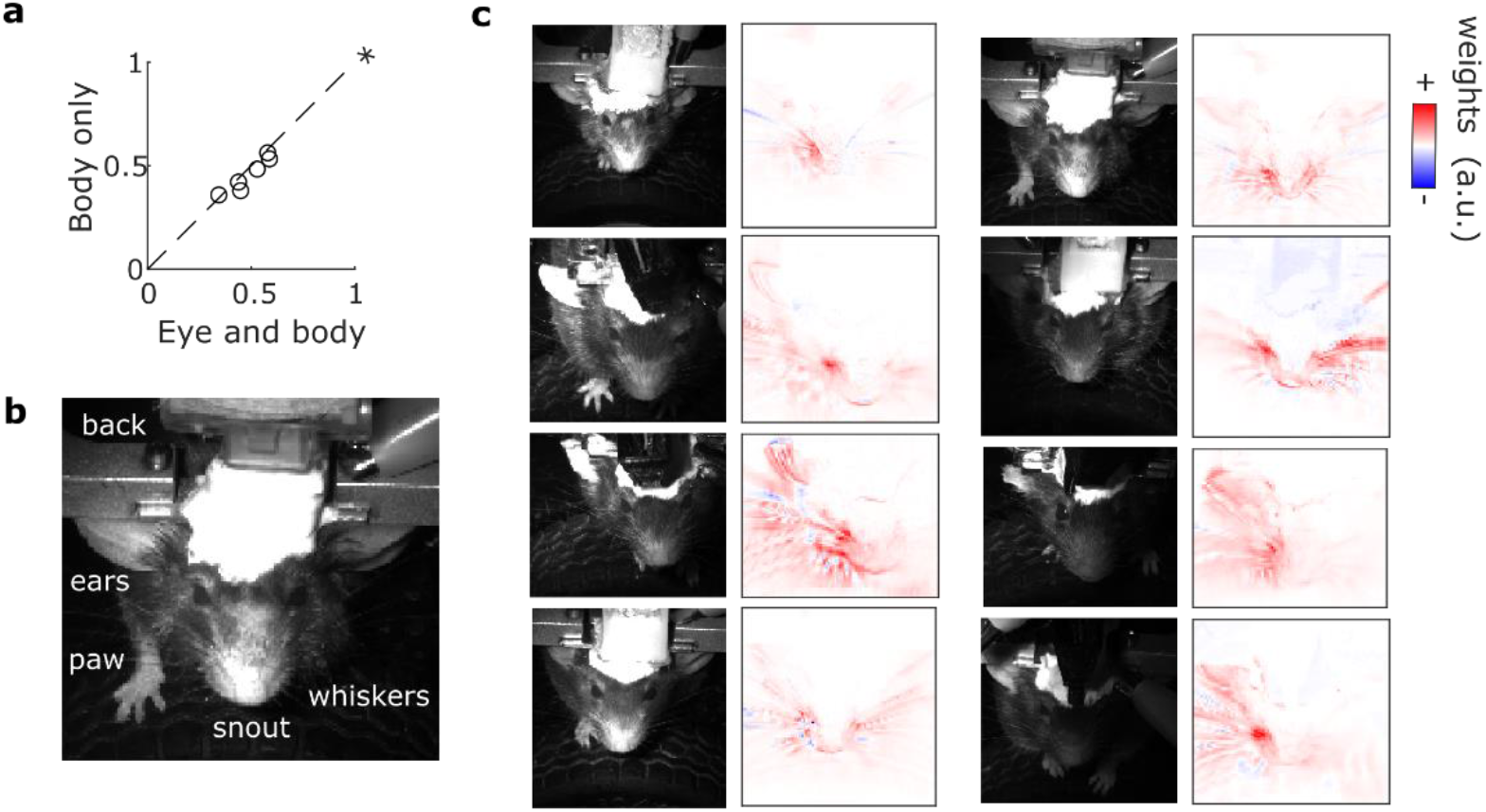
Sound-evoked V1 responses were mainly driven by whisker movements. **a**. Correlation of the actual data and their predictions for all mice, comparing different models (Eye and body: a model containing both eye and body movements predictors; Body only: a model containing body movements predictors only). The eye predictors only marginally increase the fit prediction accuracy (*: p-value<0.05), suggesting that body movements are the best and main predictors. **b**. Example frame of the face, with the different parts of the body that were visible. **c**. For each panel, left: image of a front camera recording from an example mouse; right: Reconstruction weights of the auditory PC1’s best prediction for the same mouse. Most of the weights are related to the whiskers. The asymmetry of the weight distribution across the mice’ face is mainly due to differences in lightning.

**Suppl. Figure 10.**
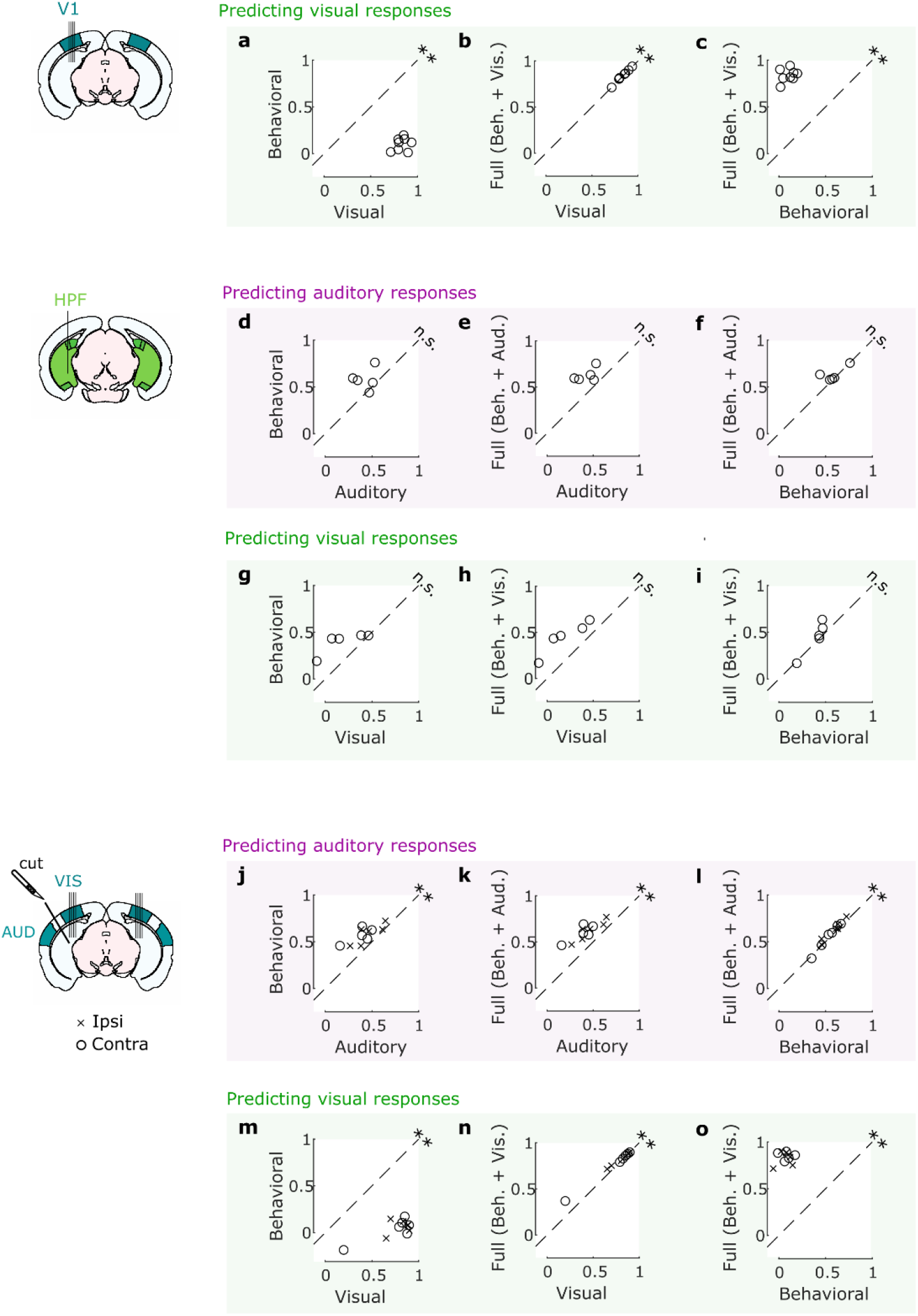
Movements explained auditory (but not visual) responses in visual cortex, and both in HPF. **a-c.** Cross-validated correlation of the visual responses and their predictions for all mice, comparing different models (Visual: videos only, Behavioral: eye and body movements only, Full: all predictors). **d-f.** Same as **a-c** but for auditory responses for the HPF recordings (albeit the low number of animals did not allow for conclusions on significance). **g-i.** Same as **a-c** but for visual responses for the HPF recordings. **j-l.** Same as **a-c** but for auditory responses for the transectomy experiment recordings. **m-o.** Same as **a-c** but for visual responses for the transectomy experiment recordings.

**Suppl. Figure 11.**
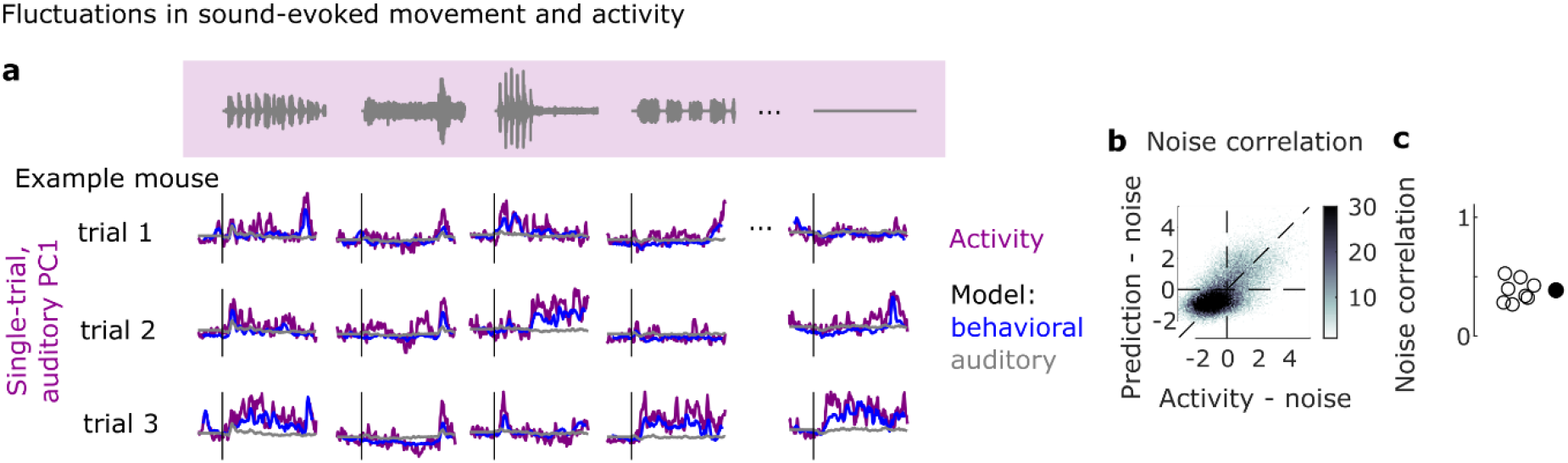
Sound-evoked movements and sound-evoked brain activity fluctuate together. **a.** Single-trial, marginalized (sound-related activity only) activity along auditory PC1 for one example mouse. The prediction from two models (auditory and behavioral) are shown. **b.** Correlation between the single-trial noise in neural activity along auditory PC1 and the single-trial noise in the prediction for an example mouse. **c.** Correlation values for all mice (empty dots) and their average (filled dot).

**Suppl. Figure 12.**
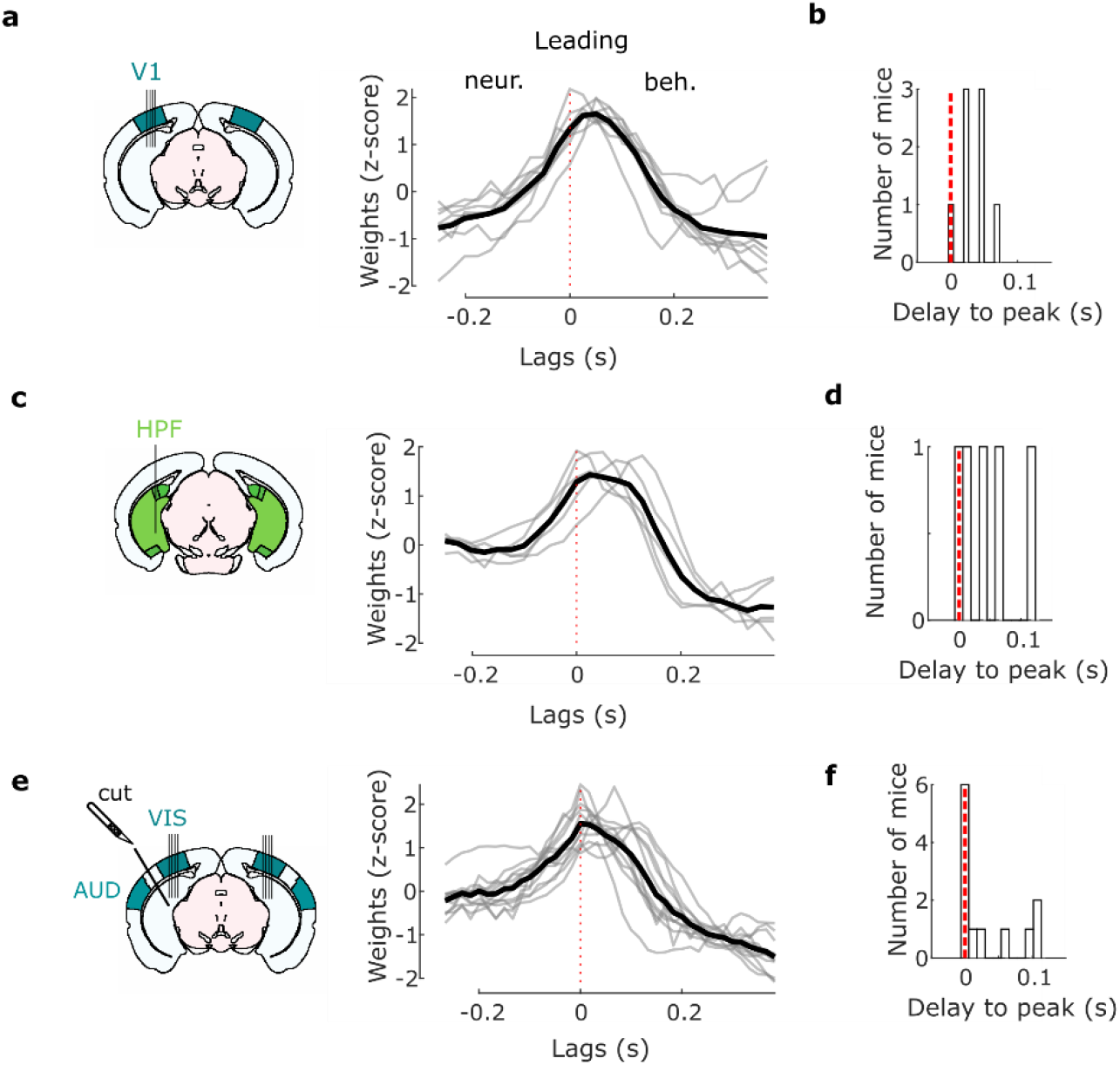
Movement precedes brain activity. **a.** Weights (motion PC1 to neural auditory PC1, z-scored) of the regression model computed on the spontaneous (no stimulus) period for the visual cortex experiments (Figure 1), showing that movement precedes neural activity (think grey lines, individual mice; thick black line: average). **b.** Distribution of the delay to the peak of the weights. A positive delay means that movement precedes and predicts neural activity by such a delay. **c, d.** Same as **a**, **b**, but for recordings in the HPF (Figure 2). **e,f.** Same as **a, b**, but for recordings in visual cortex during the transectomy experiment (Figure 3) (both sides).

